# Robust banded protoxylem pattern formation through microtubule-based directional ROP diffusion restriction

**DOI:** 10.1101/844357

**Authors:** Bas Jacobs, Jaap Molenaar, Eva E. Deinum

## Abstract

In plant vascular tissue development, different cell wall patterns are formed, offering different mechanical properties optimised for different growth stages. Critical in these patterning processes are Rho of Plants (ROP) proteins, a class of evolutionarily conserved small GTPase proteins responsible for local membrane domain formation in many organisms. While the spotted metaxylem pattern can easily be understood as a result of a Turing-style reaction-diffusion mechanism, it remains an open question how the consistent orientation of evenly spaced bands and spirals as found in protoxylem is achieved. We hypothesise that this orientation results from an interaction between ROPs and an array of transversely oriented cortical microtubules that acts as a directional diffusion barrier. Here, we explore this hypothesis using partial differential equation models with anisotropic ROP diffusion and show that a horizontal microtubule array acting as a vertical diffusion barrier to active ROP can yield a horizontally banded ROP pattern. We then study the underlying mechanism in more detail, finding that it can only orient curved pattern features but not straight lines. This implies that, once formed, banded and spiral patterns cannot be reoriented by this mechanism. Finally, we observe that ROPs and microtubules together only form ultimately static patterns if the interaction is implemented with sufficient biological realism.

## 1. Introduction

Plants are able to transport water and nutrients from the ground all the way up to the leaves, potentially more than a hundred meters high, thanks to a highly specialised system of vessels known as the xylem [1]. These xylem vessels are formed by cells that deposit a thick secondary cell wall followed by programmed cell death, leaving a hollow tube [2]. The cell wall reinforcements function to withstand the pressures generated during water transport and may be deposited in intricate patterns depending on the type of xylem. In protoxylem, the secondary wall forms bands or spirals, allowing the vessels to stretch with the surrounding tissue, while metaxylem, formed when longitudinal tissue growth has ceased, tends to be more rigid, with only some well-separated pits for radial transport [3, 4, 5] (Fig. 1A).

**Figure 1:**
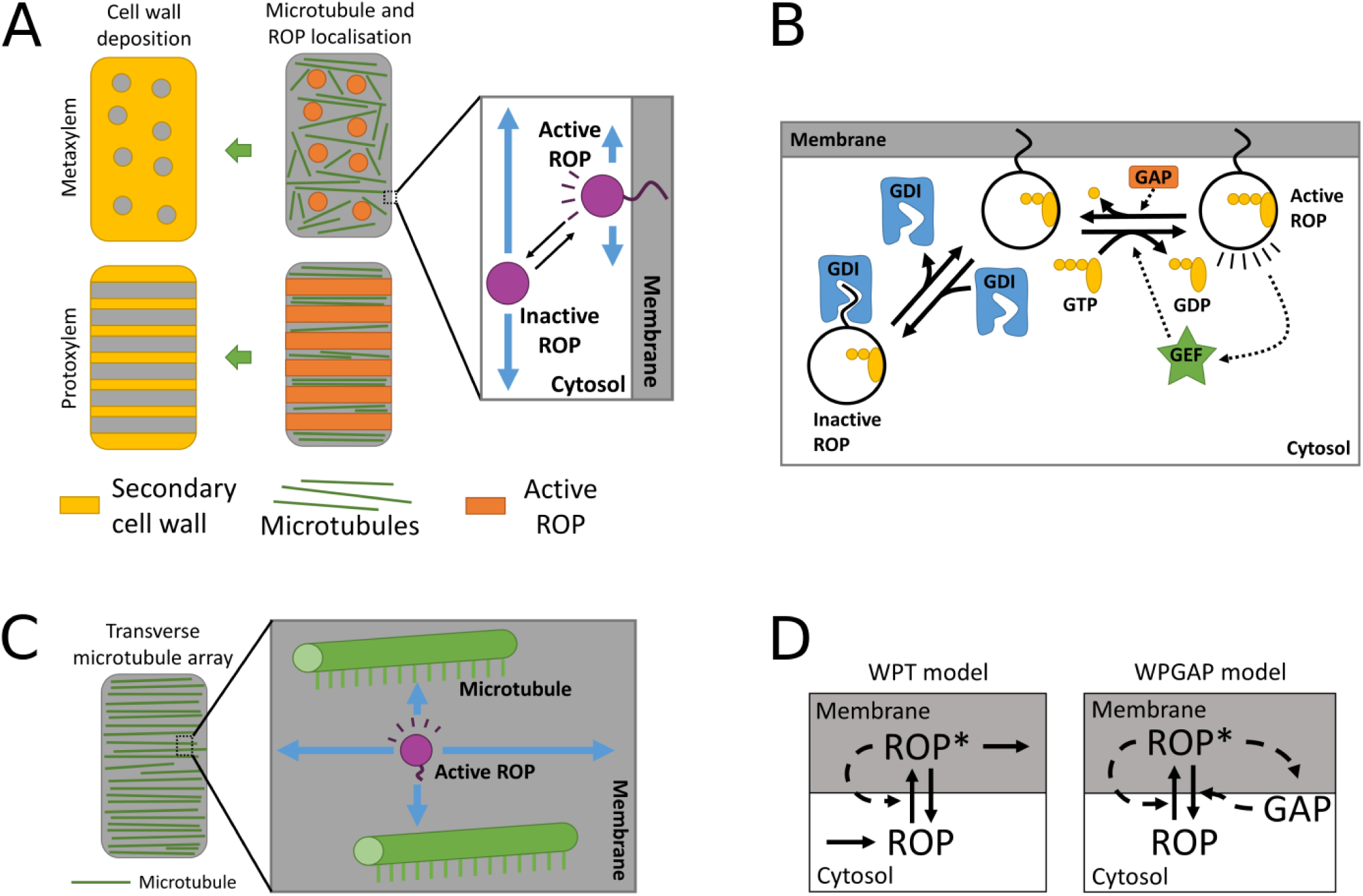
The role of ROP in xylem patterning. (A) General hypothesis of cell wall patterning in metaxylem and protoxylem. Local ROP activity destabilises the microtubules that function as a template for secondary cell wall deposition. (B) Mechanism of ROP activation and inactivation. GEFs promote ROP activation by exchanging GDP for GTP, GAPs promote inactivation by GTP hydrolysis and GDIs selectively remove inactive ROP from the membrane. (C) Cortical microtubules can self-organise into a transverse array and may act as a molecular fence, directionally restricting the diffusion of membrane-bound active ROP. (D) Interactions of active (ROP*) and inactive ROP (ROP) for two simple reaction-diffusion models for ROP-based pattern formation. Blue arrows indicate diffusion, solid black arrows indicate conversions, and dashed arrows indicate positive interactions. Panels (B) and (D) are adapted from [6].

The deposition of this secondary cell wall is determined by the position of cortical microtubules on the inside of the membrane that direct cell wall depositing cellulose synthase complexes [7, 8, 9, 10] and secretory vesicles [11]. The microtubule pattern, in turn, depends on the localised activity of ROP (Rho of Plants) proteins that recruit effectors capable of influencing microtubule dynamics, which is best demonstrated for metaxylem development. Through these effectors, ROP activity reduces the local microtubule density, creating the mirotubule pattern [12, 13, 14] (Fig. 1A). Since ROP involvement has also been indicated in protoxylem differentiation [15] and the same effectors (MIDD1, Kinesin13A) are expressed during protoxylem development [16, 17], patterning of protoxylem is expected to have a similar mechanism to that of metaxylem.

ROPs are a type of small GTPase, an evolutionarily conserved class of proteins often involved in membrane domain formation. Small GTPases are signalling proteins that function like a molecular switch. They have an active GTP-bound form and an inactive GDP-bound form (Fig. 1B). Guanine nucleotide Exchange Factors (GEFs) promote activation by exchanging GDP for GTP, while GTPase Activating Proteins (GAPs) promote inactivation by stimulating the intrinsic GTPase activity of the ROP [18, 19]. The active form is tethered to the cell membrane, while the inactive form is selectively taken out of the membrane by Guanine nucleotide Dissociation Inhibitors (GDIs) [20]. Because diffusion at the membrane is much slower than in the cytosol [21, 22], the active form diffuses only slowly compared to the inactive form. If this difference in diffusion rates is complemented by a positive feedback mechanism, all ingredients for Turing-style pattern formation are present [23, 24]. Such positive feedback loops have been demonstrated for a variety of small GTPase systems [25]. Therefore, ROPs and other small GTPases are extremely suitable for *de novo* membrane patterning. Consequently, reaction-diffusion models involving small GTPases have been proposed for a wide variety of membrane patterning processes [26, 27, 28, 29].

Generating a spotted metaxylem-like pattern using a ROP-like reaction-diffusion system is straightforward [29, 6, 30]. These same systems can also generate stripes, but these tend to be curved, without any specific orientation (e.g., patterns in first column of Fig 2), making the horizontally banded protoxylem pattern harder to obtain with a reaction-diffusion mechanism. A banded pattern can be obtained on a sufficiently narrow periodic domain, when the circumference of the domain (i.e., the width of the periodic axis) is smaller than the wave length of the pattern as suggested for, e.g., animal tails [31]. However, in microscopic pictures of protoxylem patterns, even the widest band and gap lengths combined are rarely larger than the visible width of the cell, let alone the entire circumference [10, 14]. Therefore, some other mechanism must impose the orientation of the protoxylem ROP pattern.

**Figure 2:**
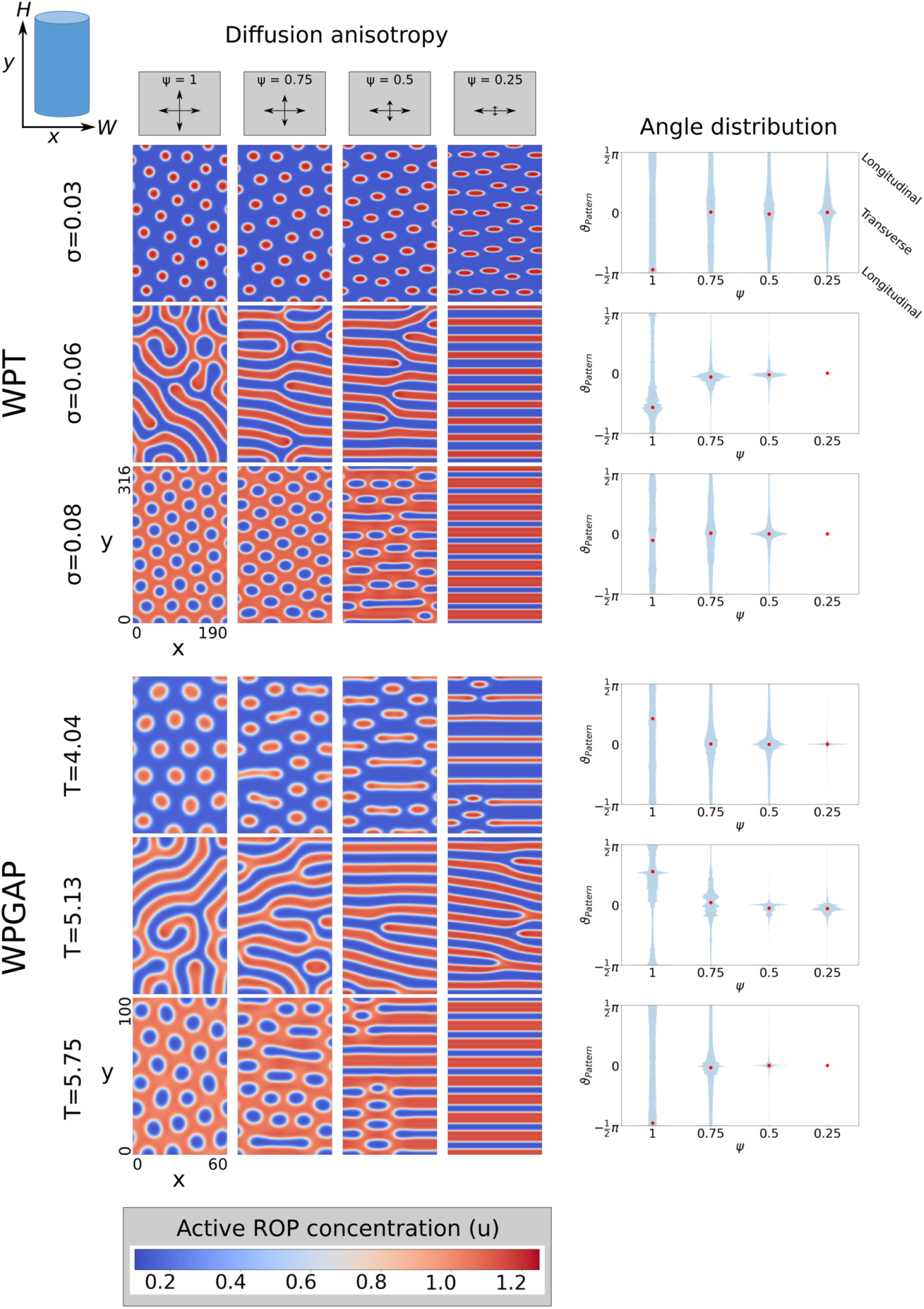
Vertical diffusion restriction of active ROP imposes a horizontal orientation on the pattern. Left: Steady state active ROP concentrations in regimes for spots, stripes, and gaps of the WPT model (top, *σ* = 0.03, 0.06, and 0.08 respectively) and the WPGAP model (bottom, *T* = 4.04, 5.13, and 5.75 respectively) for different levels of vertical diffusion restriction (*ψ*). WPT snapshots taken at *t* = 200000 and WPGAP snapshots at *t* = 40000. Right: Distributions of the pattern angles *ϑ*_*Pattern*_ (defined as orientation perpendicular to the local gradient) in the snapshots shown. Red circles indicate the average orientation. Time-lapse videos of the corresponding simulations are available online.

The protoxylem pattern has to be properly oriented relative to the cell’s growth axis, suggesting that information from the orientation of the cell is transmitted to the ROP pattern. An obvious candidate for the agent of this transmission is the microtubule array itself, since it is well-known for its ability to 90 self-organise into a wide variety of structures, including aligned cortical arrays with a transverse orientation [32, 33, 34]. This idea is consistent with the observation that, during protoxylem development, microtubules reorient to a transverse array before apparent band formation [35]. In addition, IQD13, a protein interacting with both microtubules and the plasma membrane can cause microtubules to act as a “molecular fence” that physically restricts the movement of active ROP and increased expression of this protein results in more flattened spots in metaxylem [36]. Increasing microtubule stability with taxol treatment or overexpression of MAP70 (a Microtubule-Associated Protein) has a similar effect [13, 37]. These observations suggest that, during protoxylem patterning, a transverse microtubule array will pose a barrier to any ROPs moving perpendicular to the array orientation, resulting in anisotropic ROP diffusion (Fig. 1C).

Anisotropic diffusion in reaction-diffusion systems has long been known to influence the shapes of patterns in experimental chemical systems [38] and in models used for texture synthesis [39]. A study on pattern formation in fish skin also reported pattern orientations shaped by anisotropic diffusion [40]. There, however, anisotropic diffusion was considered to be diffusion dependent on the angle of the concentration gradient, rather than a directional reduction in the diffusion coefficient. Nevertheless, since the concentration gradient determines the main direction of diffusion, their approach indirectly also results in a diffusive flow that is stronger in a certain direction. More recently, anisotropic diffusion has been proposed as a general mechanism for orienting stripes formed by Turing-like mechanisms, with horizontal diffusion restriction of a fast-diffusing component (comparable to the inactive form of ROP) resulting in horizontally oriented stripes [41]. Interestingly, in the case of protoxylem patterning we need the opposite: vertical restriction of the active form resulting in horizontally oriented stripes.

Here, we use partial differential equation (PDE) models of ROP-based reaction-diffusion systems with anisotropic diffusion to study active ROP diffusion restriction as a mechanism140 of ROP pattern orientation in protoxylem development. We also employ a functional decomposition of the diffusion tensor to gain additional mechanistic insight into the orienting power of this mechanism.

## 2. Methods

### 2.1. Modelling approach

We start with two simple reaction-diffusion models for small GTPase-based membrane patterning that can generate coexisting spots, stripes, and gaps [6]. Both models are adaptations of the wave pinning model from [42], with the addition of either protein turnover (WPT model; see Fig. 1D) or negative feedback through GAP activation (WPGAP model) to prevent accumulation of all active ROP into a single cluster (see [6] for a mechanistic explanation). The models assume that active ROP is exclusively membrane bound and inactive ROP is exclusively cytosolic. Consequently, we assume only the diffusion of active ROP will be affected by the microtubule diffusion barriers. Therefore, we will focus our investigation on the diffusion tensor *D*_*u*_ of active ROP (of concentration *u*), assuming the diffusion of inactive ROP (of concentration *v*), active GAP (*G*), and inactive GAP (*g*) to be constant and isotropic.

The dimensionless WPT model is given by:

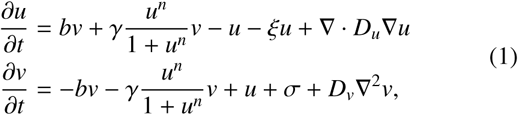

where *b* is the constant activation rate, *γ* the maximum self-activation rate, *n* the hill function exponent of self-activation, *ξ* the active ROP degradation rate, *σ* the inactive ROP production rate, *D*_*v*_ the inactive ROP diffusion coefficient, and *D*_*u*_ the dimensionless diffusion tensor for *u*. To arrive at the dimensionless forms, all diffusion coefficients and tensors were normalised with the (dimension carrying) diffusion coefficient of unrestricted active ROP (*D*_*u*,*max*_).

The dimensionless WPGAP model is given by:

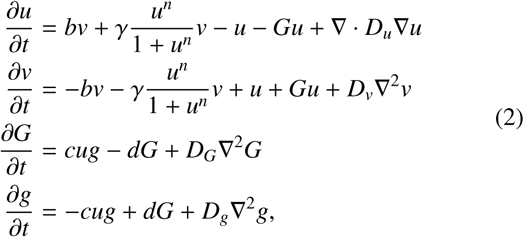

where active ROP promotes GAP activation at rate *c*, GAPs are inactivated at constant rate *d*, and *D*_*G*_ and *D*_*g*_ are the diffusion coefficients of active and inactive GAP, respectively. In the WPGAP model, the total amounts of ROP and GAP are conserved, so that the average total ROP concentration *T* and average total GAP concentration *T*_*g*_ are constants determined by initial conditions.

We used identical values for the parameters for the reaction parts and the unrestricted diffusion coefficients as in [6]. We have previously performed extensive bifurcation analyses of these models [6] to locate parameter regimes where spontaneous patterning can occur from a homogeneous state (so-called Turing regimes). This analysis directed us to relevant parameter sets that remained useful in the case of anisotropic diffusion.

### 2.2. Numerical simulations

We used an Alternating Direction Implicit (ADI) algorithm [43] to solve the reaction-diffusion equations on a two dimensional domain (see Appendix A for details on discretisation schemes). We chose a rectangular domain (190×316 or 60×100 dimensionless length units) with periodic boundary conditions in the horizontal direction and zero-flux boundary conditions in the vertical direction, representing the membrane of an elongated cylindrical cell, resembling those found in plant vascular tissue. Smaller square domains (95×95 or 63×63) with fully periodic boundary conditions were used for investigating the orientation mechanism. Unless stated otherwise, simulations were initiated at the homogeneous steady state (see [6]) with a small amount of noise added to each integration pixel. To ensure numerical accuracy, small scale tests were performed using reproducible perturbations as described by [44] to determine the optimal time step and pixel size.

### 2.3. Pattern angle analysis

To characterise the orientation of the simulated patterns, we determined the angle of the patterns *ϑ* with respect to the horizontal axis at each point on the domain. Since the direction of the pattern essentially runs perpendicular to the concentration gradient, we determined *ϑ* by taking the angle of the direction perpendicular to the gradient, such that 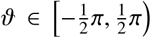. This way, *ϑ* = 0 corresponds to a transverse pattern, while 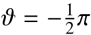 and 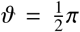 correspond to a longitudinal pattern. An average pattern angle 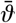 over *N* angles was calculated as follows:

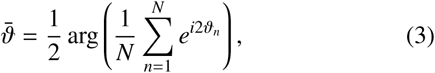

where arg is the argument or phase of a complex number and *ϑ*_*n*_ is the local angle at each integration pixel.

## 3. Results

### 3.1. Homogeneous reduction of active ROP diffusion in vertical direction

In the early stages of protoxylem patterning, before ROP activity starts to affect microtubule density, the density of the microtubule array will be more or less homogeneous. Assuming a transverse array orientation, this homogeneous vertical diffusion barrier can be represented by a spatially homogeneous reduction of active ROP diffusion in the vertical direction, yielding the following dimensionless diffusion tensor:

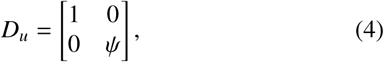

where *ψ* ∈ (0, 1] is the ratio between diffusion coefficients for restricted and unrestricted diffusion. For isotropic diffusion *ψ* = 1. Because all off-diagonal elements of *D*_*u*_ are zero, the diffusion term simplifies to:

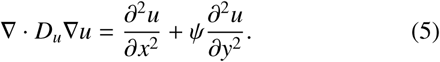

Simulation results for various values of *ψ* are shown in Fig. 2. Increasing the ROP production rate (for the WPT model) or the total amount of ROP (for the WPGAP model) changes the native pattern (i.e., the pattern formed with isotropic diffusion) from spots to stripes to gaps [6], a sequence of patterns often observed in similar models [30]. Vertical diffusion restriction of active ROP flattens spots and gaps and imposes a horizontal orientation on stripes. The stronger the restriction (the smaller *ψ*), the more the pattern is flattened. Ultimately, this turns the stripes into horizontal bands or, occasionally, spirals and forces spots and gaps to merge, resulting in banded patterns. These results show that vertical diffusion restriction of active ROP can impose a pattern of horizontal bands. Furthermore, to obtain such bands, the native pattern type does not need to be striped.

We note that the observed changes of pattern type require that the diffusion restriction only, or at least predominantly, applies to active ROP. This can be understood from a simple scaling argument: an equal diffusion reduction for both active and inactive forms is equivalent to independently scaling the x- and y-axis, which will only affect the aspect ratio of the pattern (see Appendix B for details). Similar reasoning (Appendix C) suggests that a horizontal pattern can also be obtained by horizontal diffusion restriction of the fast-diffusing component only, which has previously been demonstrated in a study comparing several other reaction-diffusion systems [41]. However, in the case of protoxylem patterning, the latter scenario is not supported by the available biological evidence and, therefore, not further investigated here.

Since both WPT and WPGAP models behave very similarly, we will continue our analyses with the WPT model only.

### 3.2. Oblique diffusion restriction promotes spiral formation

In reality, the initial microtubule array, and therefore the direction in which active ROP diffusion is restricted, does not have to be completely transverse, but may be somewhat oblique. We define *ϕ* ∈ [0, *π*) as the angle of minimal diffusion restriction with the horizontal axis. This rotates the diffusion tensor resulting in the following diffusion term for active ROP (see Appendix D for derivation):

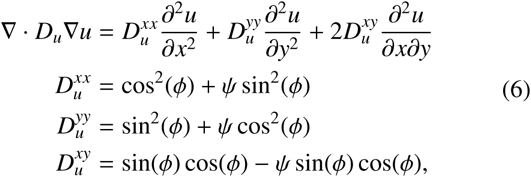

where *ψ* is again the ratio between the diffusion coefficients of active ROP in the restricted and unrestricted direction.

Such oblique diffusion restriction promotes formation of spirals (Fig. 3, Supplemental Figure S.1, where sloping lines are connected by the periodic boundary conditions in the horizontal direction). With increasing *ϕ*, double, triple, etc. spirals may form. The resulting angle of the spiral pattern *ϑ* depends on several factors. While the pattern angle depends strongly on the angle *ϕ* of diffusion restriction, particularly in early stages, it is also influenced by the boundary conditions. The periodic boundaries dictate a discrete set of preferred angles that allow connecting spirals without changing the distance between bands (Appendix E). In addition, the zero flux boundaries at the top and bottom of the domain demand that the local gradient there is perpendicular to the boundary. Consequently, spiral angles become slightly steeper than expected based on diffusion angle *ϕ* and the periodic boundaries alone.

**Figure 3:**
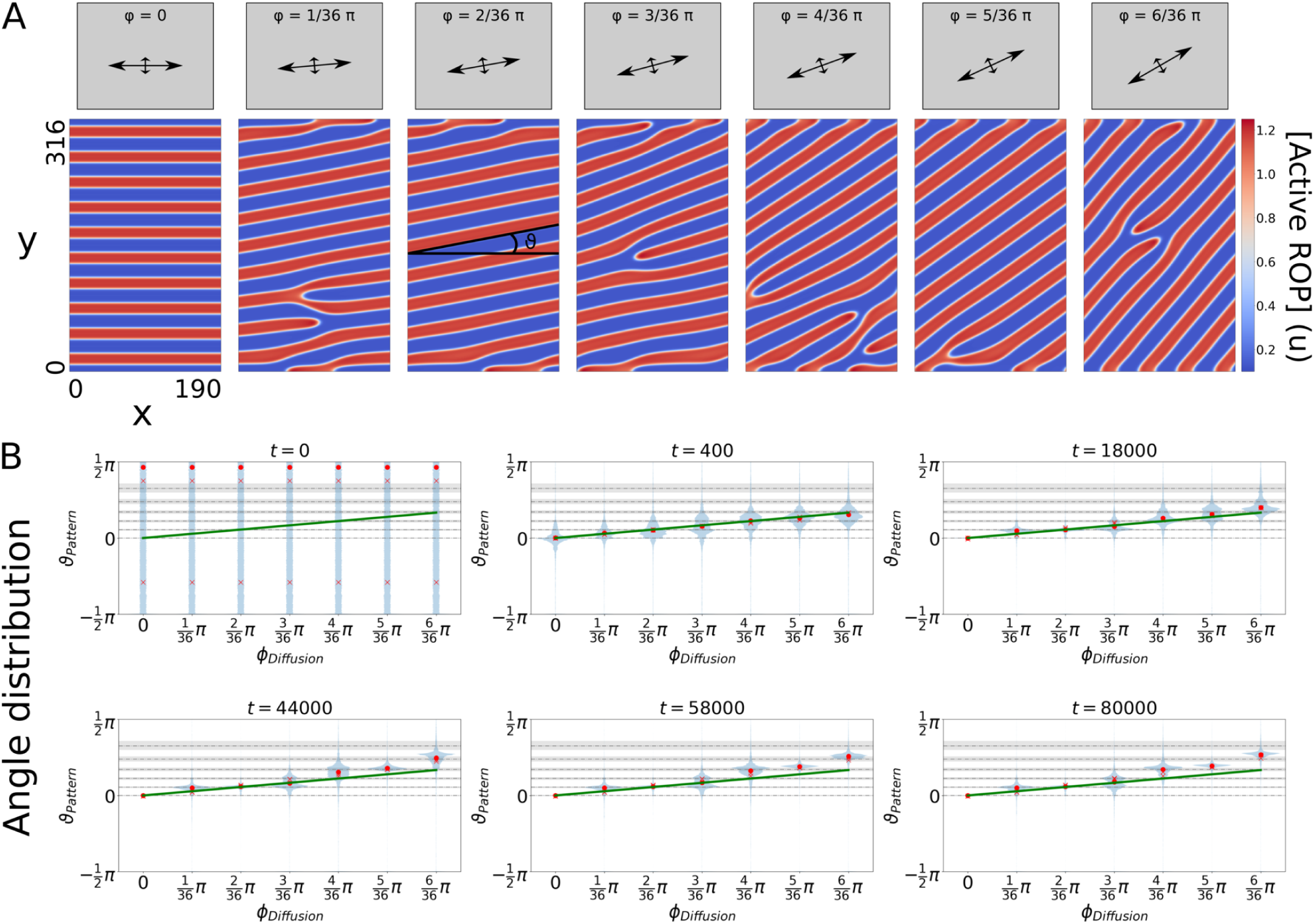
Oblique diffusion restriction of active ROP results in spirals with angles *ϑ* depending on the diffusion angle *ϕ*. (A) Active ROP concentrations at *t* = 80000 (close to steady state) in the stripe regime of the unrestricted WPT model (*σ* = 0.06) for *ψ* = 0.25. (B) Distributions of the pattern angles (defined as orientation perpendicular to the gradient) are shown for various time points. Circles indicate the average orientation, crosses indicate average orientations for replicate runs with different initial conditions (for most time points crosses are invisible due to overlap with circles), solid green lines indicate the angle of maximum diffusion *ϕ*, dashed lines indicate estimated angles for spiral patterns with band-band distances as in the banded patterns of Fig. 2, and shaded regions indicate uncertainty on those estimates due to boundary effects. Time-lapse videos of the corresponding simulations are available online.

### 3.3. Orientation mechanism and reorientability

We have seen that the restriction of active ROP diffusion in one direction promotes the formation of bands oriented perpendicular to that direction. Here, we propose a mechanism for this orientation process and investigate the implications of this mechanism on the reorientability of the pattern. Expansion of existing clusters of high ROP activity can occur when diffusion of active ROP results in ROP activation in neighbouring regions through positive feedback. Conversely, clusters of low ROP activity may expand if diffusion of active ROP from neighbouring regions lowers their activity levels such that they escape positive feedback. This suggests that the speed of expansion is related to the active ROP diffusion coefficient, so that faster diffusion in a certain direction results in faster expansion in that direction. One may hypothesise that this could lead to a change in the orientation of the pattern. From a geometrical consideration (sketched in Fig. 4A), however, it is to be expected that faster expansion in one direction can only change the orientation of curves, but not of straight lines.

**Figure 4:**
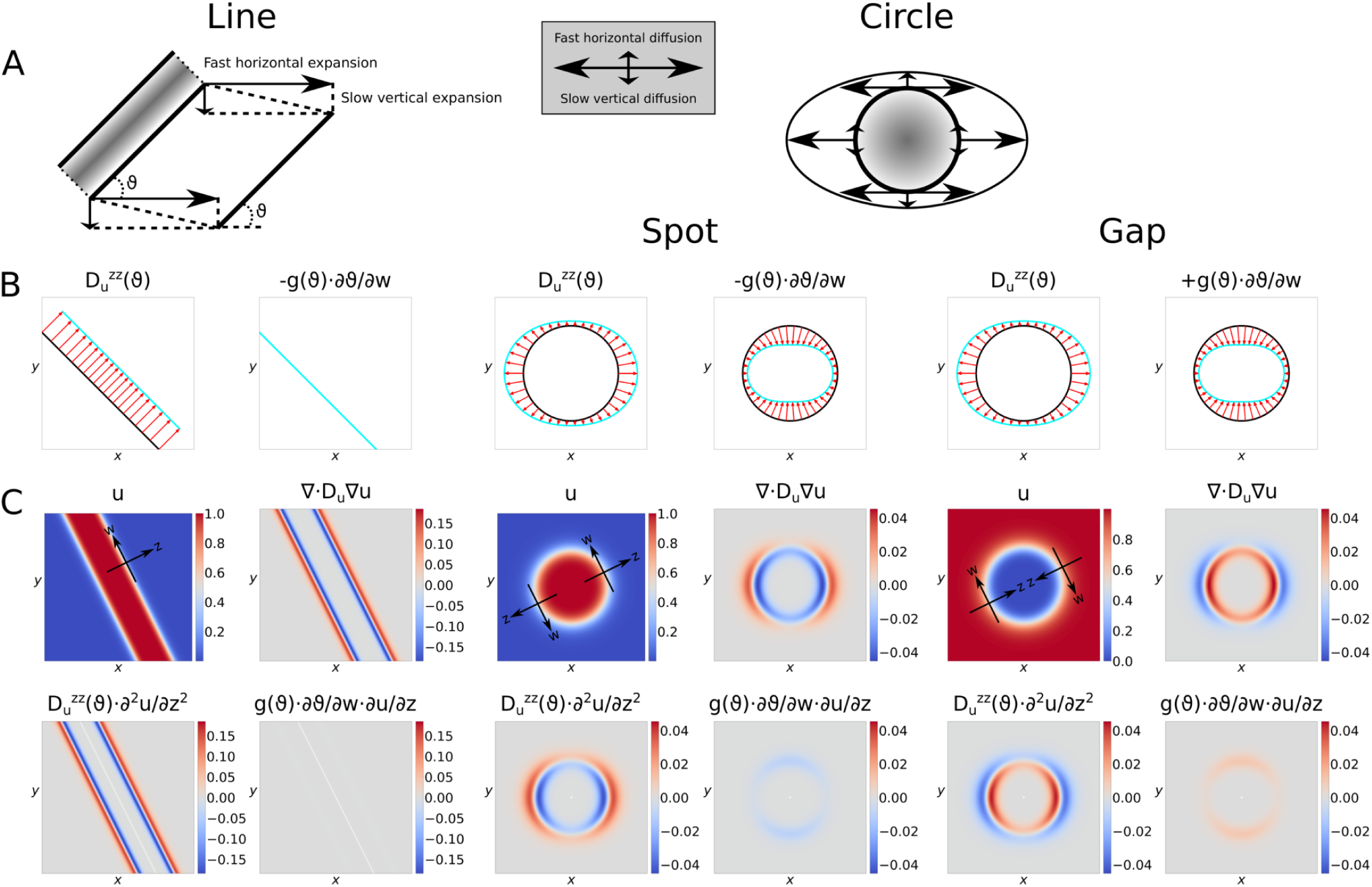
Comparison of vertical diffusion restriction effects on pattern expansion for a straight band (left) and a circular spot or gap (right). (A) Cartoon showing intuitive effect of fast horizontal expansion and slow vertical expansion. (B) Magnitude of geometry-dependent diffusion term components. Black lines indicate the local pattern geometry. Positive magnitudes are indicated by outward pointing arrows. Blue lines trace the ends of all red lines that can be drawn on the black curves. The length of the red lines reflects the magnitude of the indicated component. Since *∂u*/*∂z* is always negative and *g*(*ϑ*) is always positive, *∂ϑ*/*∂w* determines the sign of the second term from Eq. 7. To emphasise the equal magnitude but opposite effect *g*(*ϑ*) ⋅ *∂ϑ*/*∂w* has for spots and gaps, opposite signs of this term have been plotted for these two cases. (C) Magnitudes of diffusion components for stereotypical concentration profiles. Shown are: the concentration profile (*u*) with examples of axes *z* and *w*, the diffusion term corresponding to that profile (∇ · *D*_*u*_∇_*u*_), and the two separate terms of the decomposed diffusion term 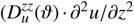 and *g*(*ϑ*) ⋅ ∂*ϑ*/∂*w* ⋅ *∂u*/*∂z*).

A decomposition of the diffusion tensor in components along and orthogonal to the pattern gradient shows that indeed only spots and richly curved patterns can be reoriented by this mechanism, but not straight lines or bands. For this decomposition, we define directions *z* and *w*, such that *z* is oriented along the concentration gradient in descending direction and *w* is perpendicular to *z* (Fig. 4C). Then we can rewrite the diffusion term from Eq. 5 in terms of *z*, *w*, and the local pattern angle *ϑ*, defined as the angle between the positive *x*-axis and the *w*-direction (see Appendix F):

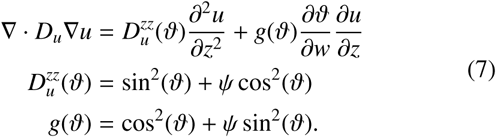

In this case, *ϑ* ∈ [−*π*, *π*). The first term resembles a standard diffusion term in the *z*-direction. Its diffusion coefficient 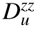 depends on *ϑ* and is therefore constant along straight lines, but larger in the unrestricted than in the restricted direction for circular patterns (Fig. 4B). The second term depends on the pattern curvature *∂ϑ*/*∂w*, which is positive for convex clusters, such as spots, and negative for concave clusters, such as gaps and zero for straight lines (Fig. 4B). Because function *g*(*ϑ*) is always positive, and *∂u*/*∂z* is always negative, the second term always represents a removal of active ROP for spots and an addition for gaps. Since *g*(*ϑ*) is largest in the restricted direction, the second term tends to promote the flattening of circular patterns.

As an illustration, we calculated the magnitude of the complete diffusion terms for some stereotypical patterns generated by the hill function:

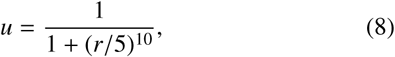

where we used *r* = 2*x* + *y* for a line and 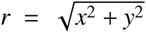 for a circle. To turn the circle from a spot into a gap we used *u*_*gap*_ = 1 − *u*_*spot*_. The results show that expansion of straight line patterns is dependent solely on diffusion down the gradient, which is the same everywhere along the line (Fig. 4C). Only curved patterns can expand in other ways, allowing their net orientation to change.

These results imply that completely non-curved patterns cannot be reoriented even if the orientation of the diffusion restriction changes, while patterns that contain curves remain open to reorientation. Simulations in which the direction of diffusion restriction is altered after straight bands have been formed confirm this (Fig. 5). Even when the native pattern consists of gaps rather than stripes, a banded pattern was maintained upon returning to isotropic diffusion (Supplemental Figure S.2).

**Figure 5:**
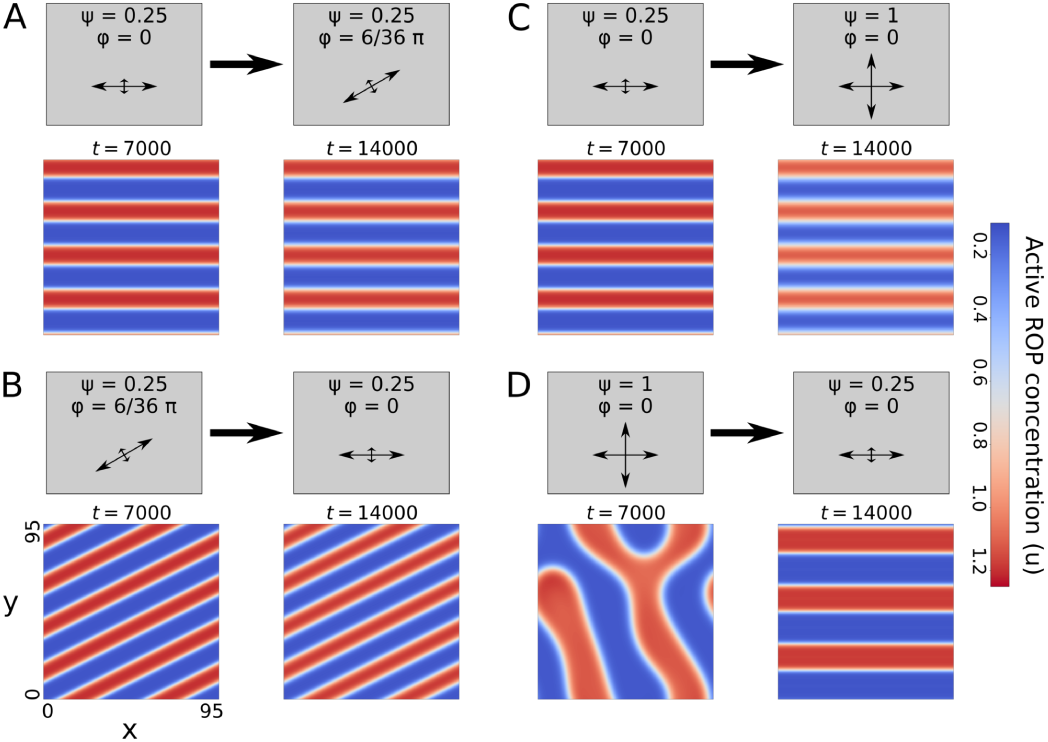
Effect of altering diffusion restriction for established patterns of the WPT model in the stripe regime (*σ* = 0.06). (A) Oblique diffusion restriction on horizontal bands. (B) Horizontal diffusion restriction on oblique bands. (C) Isotropic diffusion with horizontal bands. (D) Horizontal diffusion restriction on a pattern resulting from isotropic diffusion. Time-lapse videos of the corresponding simulations are available online.

We next investigated which kind of perturbations at the pattern level could reorient a banded pattern to better match the orientation imposed by diffusion restriction. Fully formed horizontal bands were locally stable to small amounts of spatial noise after rotating the diffusion restriction angle (Fig. 6A). Furthermore, they were also stable to moderate local vertical shifts (Fig. 6B-C), but larger shifts could trigger reorientation in the direction imposed by diffusion restriction (Fig. 6D). This reorientation is not simply an artefact of the severe deformation, since the pattern returned to the horizontal band state after applying the same deformation under vertical diffusion restriction (Fig. 6E).

**Figure 6:**
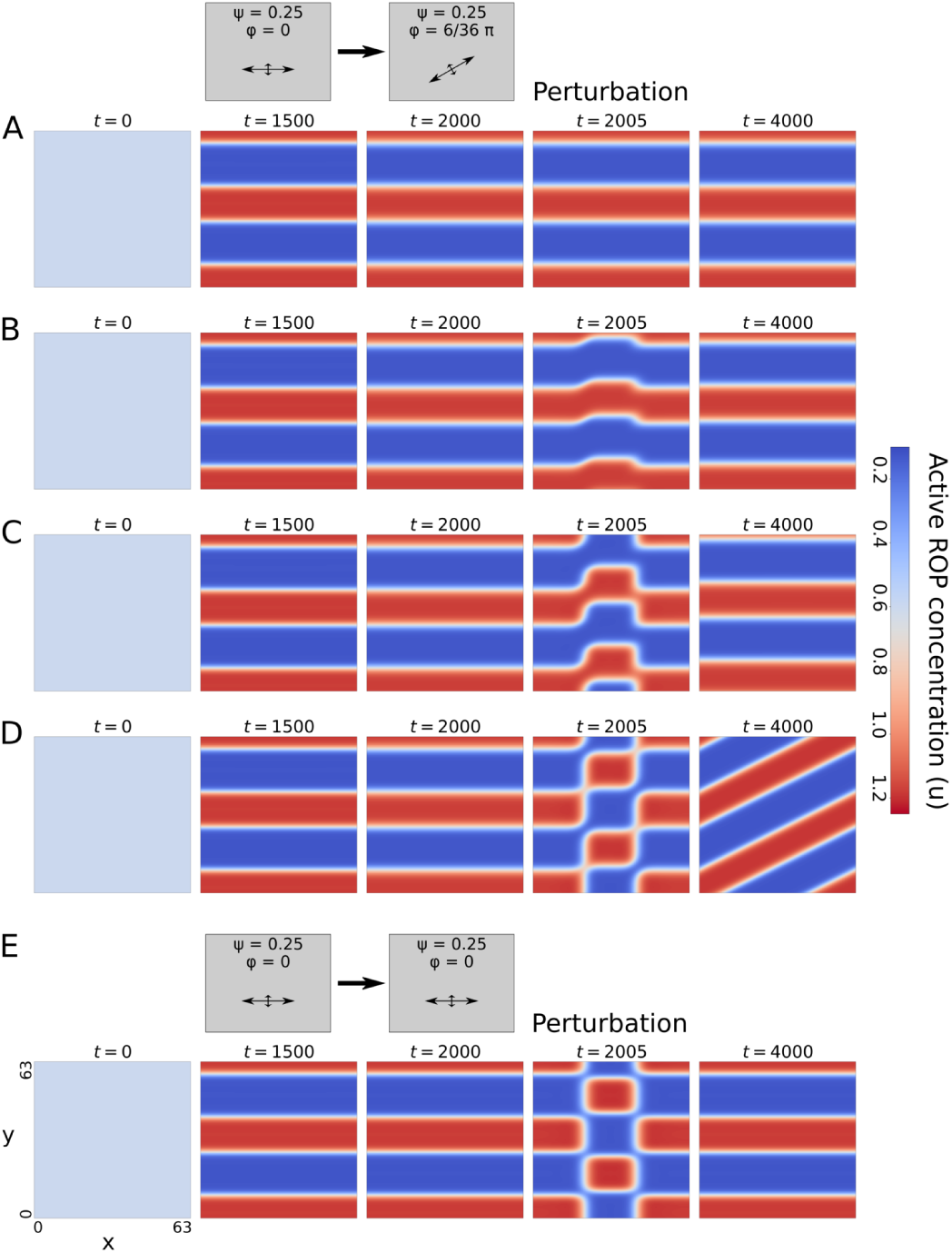
Effect of random noise and local vertical shifts on the reorientation of a horizontal band pattern with oblique diffusion restriction. A. Small random noise added every 10 time units starting from *t* = 2000. B. Small vertical shift of the central part of the pattern after *t* = 2000. C. Like B, but with a larger shift. D. Like C, but with a larger shift. E. Like D, but with vertical diffusion restriction throughout the simulation. Time-lapse videos of the corresponding simulations are available online.

These findings all suggest that diffusion restriction can impose an orientation only on patterns with curves. Since patterns start out fairly rugged from a noisy initial condition, the early stages of pattern formation are expected to be particularly susceptible to reorientation. Indeed, the preferred orientation can already be seen very early during *de novo* pattern formation (Supplemental Figure S.3). However, fully formed patterns with sufficient curvature can also still be reoriented (Fig. 5D).

### 3.4. ROP-dependent ROP diffusion

Initially, the microtubule array may form an approximately homogeneous barrier to the vertical diffusion of active ROP. Ultimately, however, ROP activity is presumed to result in gaps in the microtubule array (see Fig. 1A), which, in turn, determines the final cell wall pattern (through recruitment of downstream targets). This also means that the diffusion restriction will decrease in these gaps. To study this effect, we made the ratio *ψ* between unrestricted and restricted active ROP diffusion coefficients a function of active ROP concentration *u*, such that:

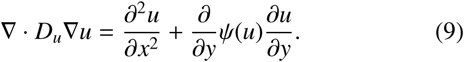

To start, we assumed a linear relation between *ψ* and the dimensionless microtubule density *ρ* (Fig. 7A):

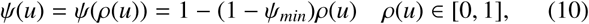

where *ψ*_*min*_ is the ratio between the maximally restricted and unrestricted active ROP diffusion components and the maximum of *ρ* is reached in absence of ROP activity. We take active ROP to reduce microtubule density instantaneously via a hill function:

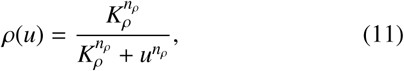

where *K*_*ρ*_ is the active ROP concentration at which microtubule density is halved and *n*_*ρ*_ is the hill exponent (see Appendix G for parameter choices). Combining Eq. 10 and 11 yields a hill function for the relation between active ROP diffusion and active ROP concentration:

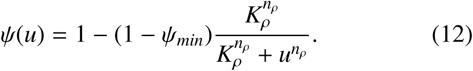

**Figure 7:**
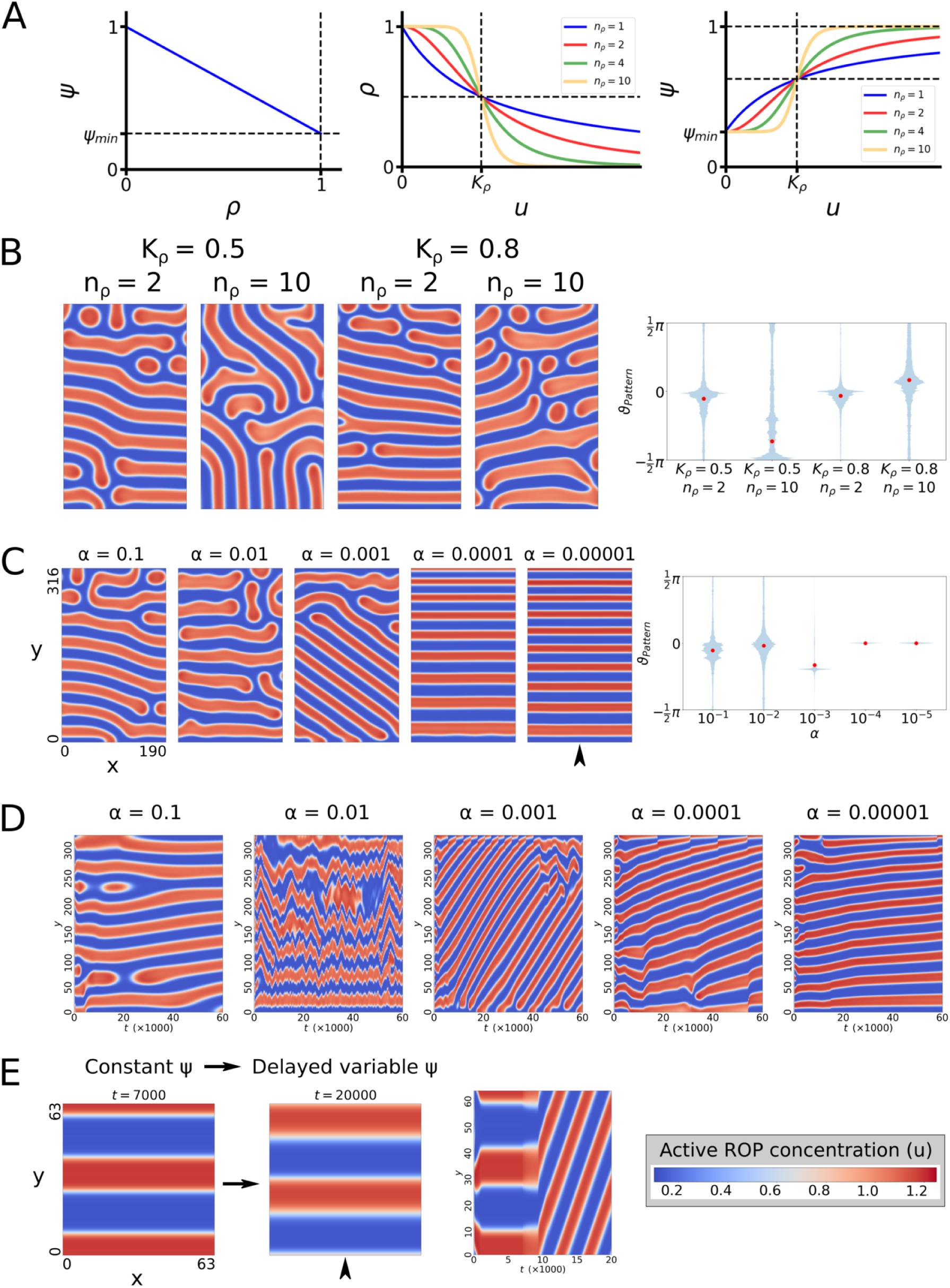
Effect of ROP-dependent microtubule density reductions on the formation of banded patterns in the stripe regime (*σ* = 0.06). (A) Relations between vertical diffusion component, (steady state) microtubule density, and active ROP concentration. (B) An instantaneous effect of ROP activity on microtubule density greatly hampers the robustness of the orientation mechanism. Snapshots at *t* = 60000. (C) Greater delays in the negative effect of active ROP on its own vertical diffusion restriction (smaller *α*) result in straighter band formation. Snapshots at *t* = 60000. (D) Time evolution of concentrations from simulations shown in (C) at the horizontal position indicated by the arrowhead. Travelling waves (recognisable by diagonal lines) occur for larger delays (smaller *α*), with wave speeds reducing as the delay becomes larger. (E) Travelling waves occur even when a static banded pattern is created first with a constant *ψ* and delayed density reduction (delayed variable *ψ*) is initiated afterwards (at *t* = 7000, using *α* = 0.001). Time evolution graph shows concentrations at the horizontal position of the arrowhead. Time-lapse videos of the corresponding simulations are available online.

Our simulations using these relations for an instantaneous effect of ROP on its own diffusion generally resulted in poorer horizontal band formation (Fig. 7B, Supplemental Figure S.4B). Some simulations got closer to horizontal bands than others, but we could no longer find a broad parameter regime that consistently yielded proper band formation, suggesting that the robustness of the orientation mechanism was greatly reduced.

### 3.5. Delayed microtubule density reduction

We saw earlier that developing patterns are particularly susceptible to reorientation, while fully formed banded patterns can be quite resistant to it. These results suggest that a more or less homogeneous diffusion restriction may only be necessary during the development of the ROP pattern, allowing patterning of the microtubule array afterwards, without influencing the ROP pattern. Therefore, we investigated the effect of a delayed response of the microtubule density to ROP activity. Taking a first order approximation for the rate of change towards steady state density *ρ*\*, we get:

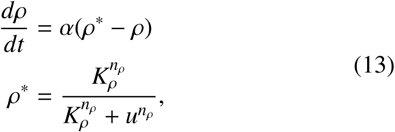

where *α* is a rate constant. This means that the diffusion ratio *ψ* will approach its steady state *ψ*\* according to:

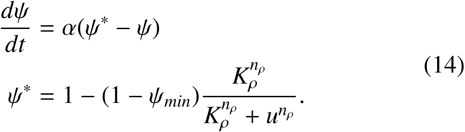

Simulations with a homogeneous starting density of *ρ* = 1 (the maximum, so *ψ* = *ψ*_*min*_) and small delays (*α* = 0.1) yielded patterns comparable to those of the no delay scenario (Fig. 7C, Supplemental Figure S.4C). Larger delays did result in straighter bands, but these bands were constantly moving, either up and down, or in one direction like a travelling wave (Fig. 7D, Supplemental Figure S.4D). These waves occurred even when starting from a fully formed static banded pattern (Fig. 7E, Supplemental Figure S.4E). Such non-static patterns are unrealistic as a basis of cell wall pattern formation.

### 3.6. Positive feedback in microtubule density can prevent travelling waves, stabilising a banded pattern

Thus far, we assumed microtubule density to be merely a reflection of ROP activity, possibly with a delay. However, microtubules have their own complex dynamics that may well have a stabilising influence on the patterning process. Therefore, we will consider a potential positive feedback loop acting on the microtubule density. Microtubule growth occurs only at the tips of existing microtubules. In addition, nucleation of new microtubules occurs predominantly from existing microtubules, with predominantly congruent orientation [45, 46, 47]. Therefore, we can expect stronger increases in microtubule density at places with higher existing density. The resulting positive feedback loop between microtubule density and microtubule nucleation might serve to stabilise local regions of high microtubule density, thereby preventing travelling wave patterns.

To test this idea, we model this positive feedback with a hill function and let ROP activity promote microtubule degradation, so that dimensionless microtubule density *ρ* obeys the equation:

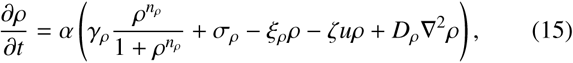

where *γ*_*ρ*_ is the maximum microtubule-dependent growth rate, *n*_*ρ*_ is the hill exponent, *σ*_*ρ*_ is a constant background increase in microtubule density by nucleation and growth, *ξ*_*ρ*_ is a constant linear degradation rate, and *ζ* is a ROP-dependent degradation rate. Here, *ρ* is scaled with the density at which positive feedback is half its maximum. In addition, we include a diffusion term with coefficient *D*_*ρ*_ to maintain a connection between neighbouring integration pixels and prevent artefacts. We take *D*_*ρ*_ ≪ 1 to keep microtubule “diffusion” well below unrestricted active ROP diffusion. Although microtubules don’t really diffuse in their entirety, they tend to spread through growth and nucleation. These processes also favour the existing orientation [46], which could further promote stable bands. Here, we assume isotropic diffusion as a worst case scenario. Finally, we use α as a tuning parameter for the time scale of the microtubule dynamics relative to the ROP dynamics.

The steady state of the homogeneous system is determined by:

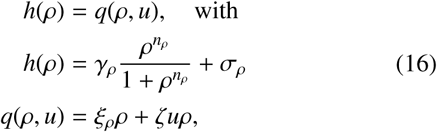

which allows for bistability as long as *n*_*ρ*_ > 1 (Fig. 8A). In that case, the steady state microtubule density may switch from low to high or the other way around as ROP activity changes.

**Figure 8:**
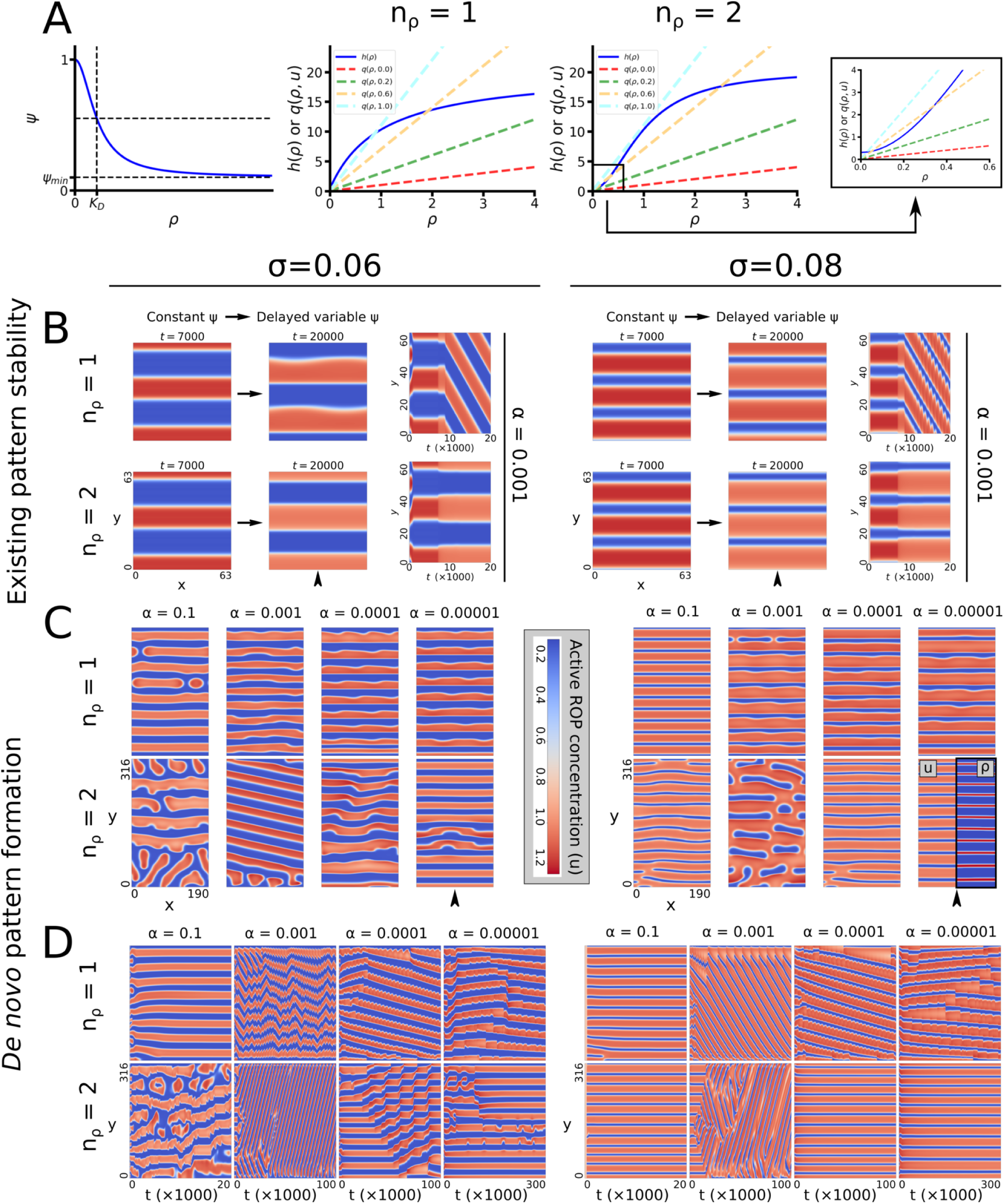
Positive feedback in microtubule density can prevent bands from moving vertically. All simulations were performed in both the stripe (*σ* = 0.06) and the gap (*σ* = 0.08) regime of the WPT model. (A) Left: Relation between microtubule density (*ρ*) and vertical diffusion of active ROP. Right: Values of hill function and linear components of the microtubule density equation for various values of *u*. Intersections indicate steady states. Bistability can occur only for *n*_*ρ*_ > 1. (B) Stability of a banded pattern established with a constant microtubule density until *t* = 7000 for unistable (*n*_*ρ*_ = 1) and bistable (*n*_*ρ*_ = 2) microtubule dynamics. In all cases, *α* = 0.001 and fully periodic boundary conditions were used. Graphs to the right show time evolution of concentrations at the horizontal position of the arrowhead. (C) Full *de novo* patterning with positive feedback of microtubule density starting from the homogeneous state for ROP concentrations and *ρ*_0_ = 2. Snapshots were taken at the final time points indicated in (D). Inset in bottom right snapshot shows microtubule densities (*ρ*) for comparison. (D) Time evolution of simulations from (C) at the horizontal position indicated by the arrowhead. Time-lapse videos of the corresponding simulations are available online. Corresponding microtubule densities (*ρ*) for the entire figure are shown in Supplemental Figure S.5.

We use a hill function for ψ to ensure that vertical diffusion is close to its maximum (*ψ* = 1) for low densities, and close to its minimum (*ψ* = *ψ*_*min*_) for high densities (Fig. 8A):

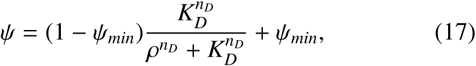

where *K*_*D*_ is the density at which vertical diffusion is halfway between its minimum and its maximum and *n*_*D*_ is a hill exponent.

Even when starting from a banded pattern, the positive feedback model without bistability (*n*_*ρ*_ = 1) at moderate delay (*α* = 0.001) yielded travelling waves (Fig. 8B), just like we observed for a direct ROP-dependency with a simple delay. However, when we included bistability (*n*_*ρ*_ = 2), the banded pattern remained static.

However, when simulating *de novo* patterning of a large domain we still observed non-static patterns at *α* = 0.001, even with bistability (Fig. 8C–D). For large delays (*α* = 0.00001), a banded ROP pattern had sufficient time to develop before significant changes in microtubule density occurred and the banded pattern remained stable. Interestingly, a stable banded pattern seems to form more readily in the gap regime (*σ* = 0.08) than in the stripe regime (*σ* = 0.06). We also found some static banded patterns, particularly in the gap regime, for small delays (*α* = 0.1) that were apparently insufficient to trigger travelling waves.

For the static patterns, microtubule density is essentially a readout of ROP activity (Fig. 8C, inset). For the moving patterns, especially in the bistable case, the delay causes the microtubule density to lag behind the ROP pattern and never fully reach its high steady state (Supplemental Figure S.5), as the pattern has already moved on before it got the chance. This may also explain the stuttering of the moving patterns in the bistable case for ROPs and microtubules.

These results show that the precise implementation of microtubule dynamics and the interactions between ROPs and microtubules matters and inclusion of more realistic biological details seems to result in more stable banded patterns.

## 4. Discussion

### 4.1. Directional diffusion restriction can orient ROP patterns into bands and spirals

We have shown that restricting active ROP diffusion in a specific direction is sufficient to change the output from a ROP-based Turing-style pattern formation mechanism from patterns without specific orientation into banded or spiral patterns with controlled orientation. Given sufficiently large diffusion restriction, bands and spirals can be generated, regardless of whether the native pattern would consist of spots, stripes, or gaps. The parameters that normally distinguish between these regimes then control only the thickness of the bands. This means that the relatively thick active ROP bands that would be needed to generate the relatively thin microtubule bands observed in protoxylem patterning [10] may actually originate from a Turing mech-445 anism that would generate gaps without diffusion restriction. Additionally, the finding that diffusion restriction can essentially turn spot, stripe, and gap regimes into one large band regime may imply an increased robustness of the protoxylem pattern to changes in ROP activity. By comparison, the spotted metaxylem pattern would more easily be disrupted.

At intermediate levels of diffusion restriction, spots, stripes, and gaps can still be distinguished, but they are flattened horizontally. This finding is in line with the experimental observation that higher expression of a protein responsible for a stronger microtubule barrier effect results in more flattened pits in metaxylem [36]. Therefore, protoxylem patterning may be very similar to metaxylem patterning, with a stronger diffusion barrier effect to obtain a banded pattern and a higher ROP expression to obtain thicker ROP bands. In addition, our finding that the angle of diffusion restriction largely determines the angle of *de novo* banded pattern formation, predicts that the initial orientation of the microtubule array before apparent band formation determines the final orientation of the banded array.

While the precise molecular players and interactions involved in protoxylem patterning remain largely unknown, ROP involvement has been implicated [15]. In addition, we show that different ROP-based Turing mechanisms have a very similar response to a directional reduction in activator diffusion. Therefore, the proposed orientation mechanism seems sufficiently general to apply to different reaction-diffusion systems able to generate coexisting spots, stripes, and gaps, of which many variants exist [6, 30, 29]. However, it is important that diffusion restriction applies predominantly to the active form, because diffusion restriction of all components in the same way will not yield banded patterns with controlled orientation (Appendix B). The inactive form must therefore be able to bypass the microtubule barriers. This requirement is naturally fulfilled by the predominantly cytosolic localisation of the inactive form.

### 4.2. Fully formed banded patterns are relatively robust structures compared to more curved patterns

While anisotropic diffusion has long been known to affect the shape of patterns generated by reaction-diffusion systems [38, 39], we not only demonstrated its potential for generating bands and spirals, but also investigated the underlying orientation mechanism. Our decomposition analysis of the diffusion term suggests that if ROP cluster expansion is related to diffusion of the active form, any curved structures tend to expand more in the direction in which diffusion is unrestricted. However, straight patterns cannot be reoriented in this way, making fully formed bands and spirals insensitive to any subsequent changes in the direction of diffusion restriction, even in the presence of moderate spatial perturbations.

This robustness of straight patterns has several benefits for the biological system. Firstly, in order to create the actual cell wall pattern, the underlying ROP and microtubule patterns will need to remain approximately stationary for the duration of secondary cell wall synthesis, which may take many hours [10], so the ROP pattern should be robust to any perturbations that may occur in this time frame. Secondly, due to the difficulty of reorienting a pattern of oblique bands, it seems important that the reorientation of the microtubule array to a transverse state occurs before the start of ROP patterning. Experimental observations suggest that this reorientation at least occurs before visible band formation in the microtubule array [35]. In addition, aberrant protoxylem patterns have been observed in the Katanin mutant, where proper microtubule alignment prior to band formation is hindered [48]. Finally, once a transverse microtubule array has contributed to the generation of a banded ROP pattern, the robustness of the banded pattern allows microtubules to disappear in areas of high ROP activity without disrupting the ROP pattern.

### 4.3. Static band formation depends on the precise implementation of microtubule dynamics

Our findings show that an instantaneous or very fast response of microtubule density to ROP activity can be disruptive to proper band formation. A sufficiently slow microtubule response, however, gives a robust banded pattern time to form before the diffusion restriction is undone. However, this dynamic also results in a kind of delayed negative feedback, which is known for its potential to generate oscillations [49] and travelling waves [50, 51] in models. With more realistic microtubule dynamics, we were able to stabilise the banded pattern again. The required difference in time scales of ROP protein and microtubule density dynamics (approximately a factor *α*) does, however, seem quite large, particularly where the native pattern would consist of stripes. Partly, a large difference may actually be realistic, since changes in microtubule density during protoxylem patterning can take hours [10], whereas GTPase inactivation and removal from the membrane occurs in the order of seconds [52, 53]. Also, since the microtubule bands formed in protoxylem patterning are relatively thin [10], the underlying ROP pattern must have relatively thick bands, like those from the gap regime, in which generation of static bands occurred already with smaller delays.

Other biological factors not included here may also help stabilise a banded pattern, potentially reducing the required difference in time scales. Firstly, ROP pattern formation may already start before the ROP effectors that interact with microtubules become active, allowing the ROP pattern time to establish without ROP activity having any impact on microtubule density. Transcriptomic data seem to suggest that, in metaxylem, expression of effectors like MIDD1 and Kinesin-13A peaks later than that of ROP11, although the temporal resolution is not very good [54]. Secondly, microtubules are not simply slowly diffusing molecules, but discrete linear structures with complex dynamics and membrane attachment preventing translational movement. Their rigid and linear nature may well contribute to the formation of robust linear bands. Furthermore, microtubule-dependent microtubule nucleation tends to favour the direction of the parent microtubule [46], so that any partial microtubule bands will tend to expand in the direction they already have. Finally, microtubule dynamics are governed largely by processes occurring at their growing or shrinking tips [55]. This suggests ROP activity cannot destabilise existing microtubules except at their ends. Such details of microtubule dynamics have no straightforward implementation in the present modelling paradigm, but detailed stochastic simulation models of the cortical microtubule array with explicit dynamic instability exist [56, 57, 58, 59, 60]. Building upon those, a hybrid model combining PDE integration for ROP dynamics with stochastic microtubule dynamics offers an exciting direction for future research on protoxylem patterning.

**Table 1:** Parameter values used for ROP-dependent ROP diffusion. ROP can depend on the active ROP concentration either instantaneously (Instant), or with a simple delay (Delayed), or with a microtubule density relation including positive feedback (MT positive feedback). For empty fields that version of the model does not include a parameter with that name.

| Parameter | ROP-dependent ROP diffusion relation |  |  |
| --- | --- | --- | --- |
|  | Instant | Delayed | MT positive feedback |
| $\psi_{min}$ | 0.25 | 0.25 | 0.1 |
| $n_\rho$ | Variable | 2 | 1 or 2 |
| $K_\rho$ | Variable | 0.5 | |
| $\alpha$ | | Variable | Variable |
| $\rho_0$ | | 1 | 2 |
| $\gamma_\rho$ | | | 20 |
| $\sigma_\rho$ | | | 0.3 |
| $\xi_\rho$ | | | 1 |
| $\zeta$ | | | 10 |
| $n_D$ | | | 2 |
| $K_D$ | | | 1 |
| $D_\rho$ | | | 0.01 |

## Acknowledgements

We thank Tijs Ketelaar for insightful discussions on the topic.

## Appendices

## A. Discretisation of the diffusion equation for numerical integration

In general, a diffusion term has the following form:

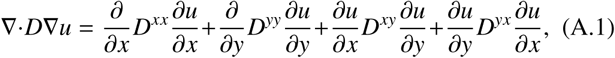

where *u* is the concentration of the diffusing component, D is the diffusion tensor, and *D*^*xy*^ = *D*^*yx*^. When the diffusion coefficient is homogeneous (i.e. independent of space), this simplifies to:

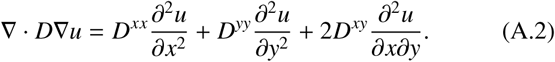

In this case, the diffusion coefficients do not need to be discretised and discretising the concentrations is straightforward, with *u*_*xx*_ at position (*x*_*i*_, *y*_*j*_) given by:

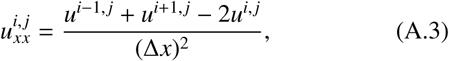

where ∆*x* is the spatial step size in the x-direction. The discretisation of *u*_*yy*_ is done in the same way. For cases with oblique diffusion restriction, *u*_*xy*_ can be discretised as follows:

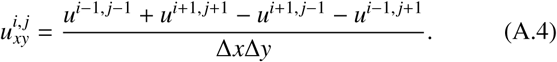

For periodic boundary conditions, the last grid point in the periodic direction is simply treated as if it were attached to the first grid point in that direction. For zero-flux boundary conditions, an extra virtual grid point is added at the ends with the same concentration as the grid point before, such that:

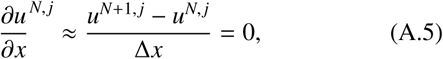

or the equivalent in the y-direction, where *N* is the number of the last grid point. As long as at least directions has periodic boundary conditions, this scheme will guarantee mass conservation.

For inhomogeneous diffusion, the diffusion coefficients from Eq. A.1 also need to be discretised, because they depend on *x* and *y*. For the diffusion term in the x-direction, this was done as follows:

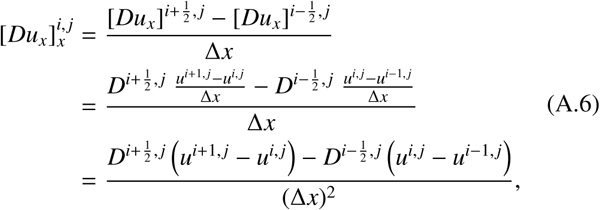

where *D* represents *D*^*xx*^. The diffusion term in the y-direction can be discretised in the same way. This scheme requires the concentrations to be known at the grid points and the diffusion coefficients between grid points (Fig. A.1). To determine the diffusion coefficients between grid points, the average of the two nearest neighbouring grid points was used. For periodic boundary conditions, an extra diffusion node can be added, connecting the last and the first grid points. Zero-flux boundary conditions can be imposed in the same way as before, or by setting the diffusion coefficient between the last grid point and the virtual grid point equal to zero.

**Figure A.1:**
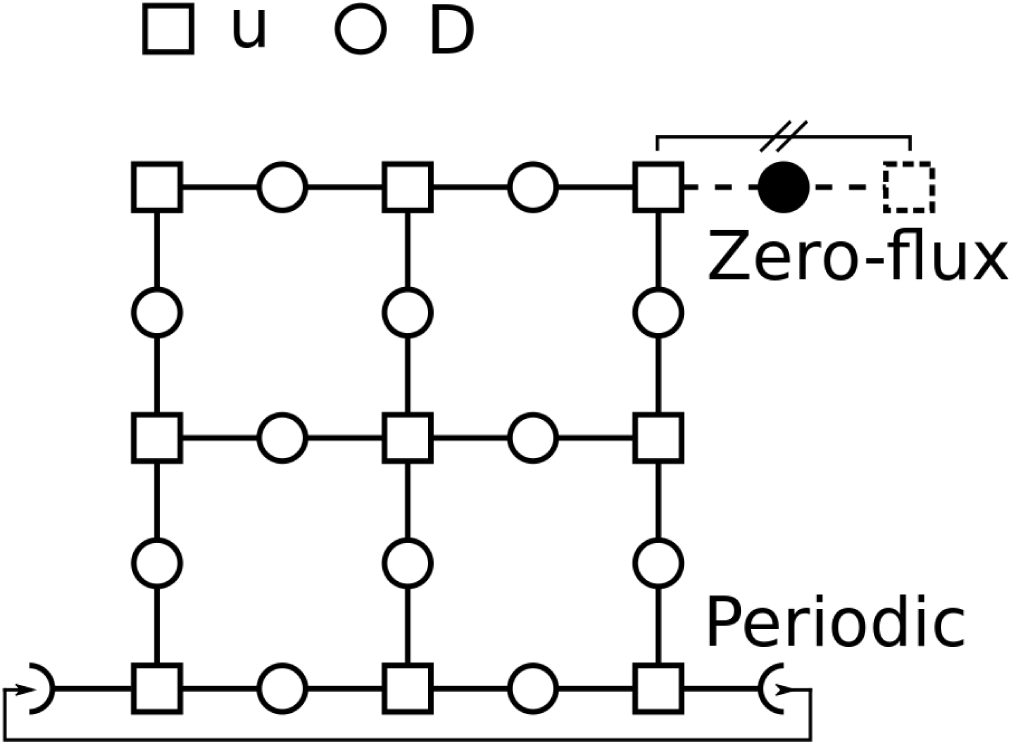
Discretisation scheme and boundary conditions for models with inhomogeneous diffusion coefficients. Squares indicate grid points at which concentrations (*u*) are known. Open circles indicate points at which diffusion coefficients are imputed. Closed circles indicate diffusion coefficients set to zero for zero-flux boundary conditions.

If oblique diffusion restriction were to be used as well, the *xy*-terms would also need to be discretised. This can be done as follows:

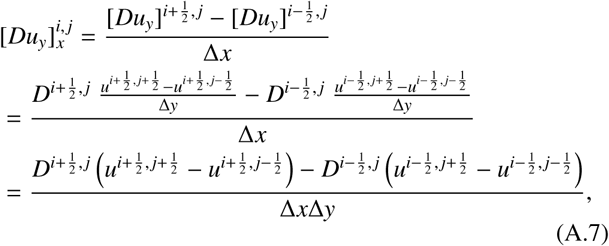

where *D* represents *D*^*xy*^. The other *xy*-term can be discretised in the same way. This scheme also requires concentrations at half steps in both directions (Fig. A.2). These can be calculated by taking the average of the four nearest neighbours. For periodic boundary conditions these averages also have to be determined between the last and first grid points. For zero-flux boundary conditions, additional virtual concentration points are required that have the same value as their neighbours. Again, these schemes will guarantee mass conservation as long as at least one of the directions has periodic boundary conditions.

**Figure A.2:**
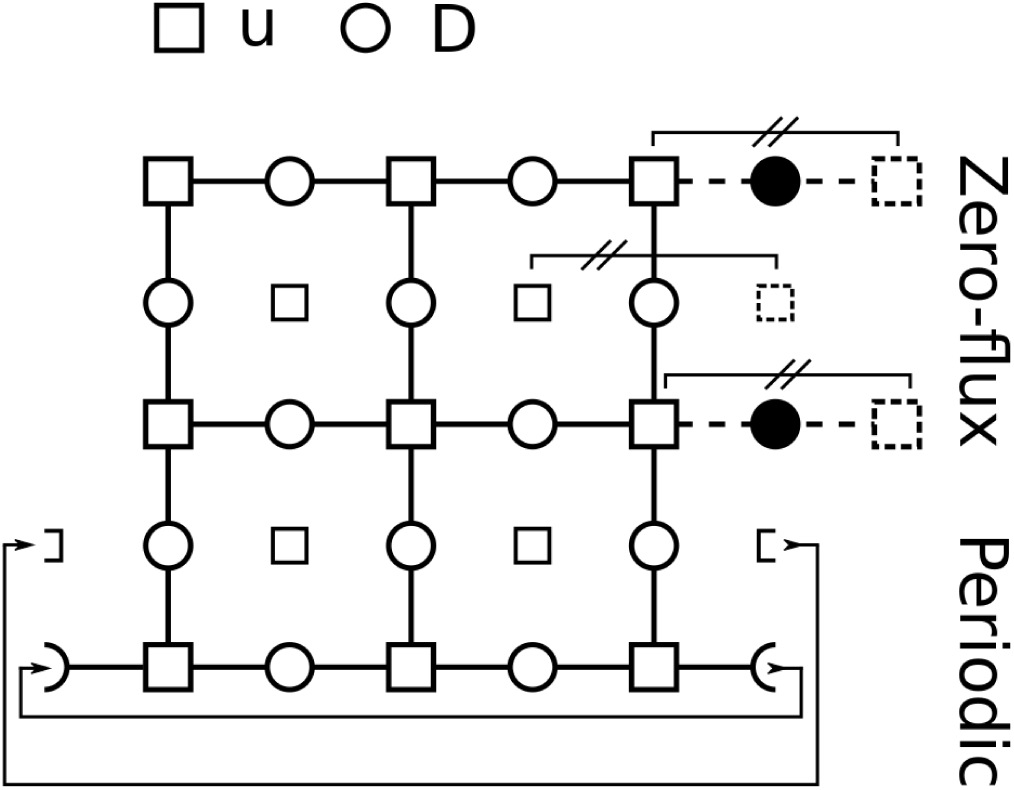
Discretisation scheme and boundary conditions for the *xy*-terms of models with inhomogeneous diffusion coefficients and oblique diffusion restriction. Large squares indicate grid points at which concentrations (*u*) are known. Smaller squares indicate grid points at which the concentrations need to be imputed. Open circles indicate points at which diffusion coefficients are imputed. Closed circles indicate diffusion coefficients set to zero for zero-flux boundary conditions.

## B. Effect of vertical diffusion reduction for all variables

In this study, we have focused on vertical diffusion restriction of active ROP only, which is the scenario that seems most congruent with the biological evidence. Here, we will elaborate a simple scaling argument for why vertical diffusion restriction of all components, in fact, cannot similarly function as a mechanism for obtaining horizontally banded patterns.

For every variable w in both the WPT and WPGAP model, we have:

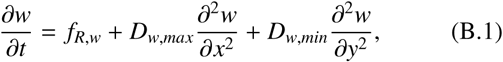

where *f*_*R*,*w*_ is the reaction function of *w* and other variables. If the vertical diffusion coefficient is reduced by a factor *q*, so that:

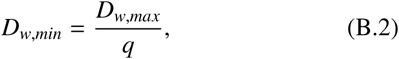

then we get:

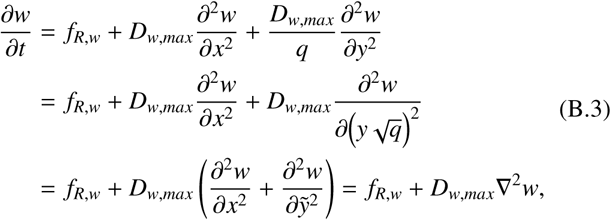

where 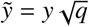. This shows that reducing the vertical diffusion coefficients of all variables by a factor *q* yields the same results as compressing vertical space by a factor 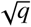.

## C. Effect of alternative anisotropic diffusion scenarios

This is a purely mathematical consideration of a alternative scenarios for obtaining banded patterns using anisotropic diffusion. There are no biological indications supporting these scenarios in the case of xylogenesis.

In our main scenario of vertical restriction of active ROP diffusion, only the equation for *u* can be scaled as in Appendix B:

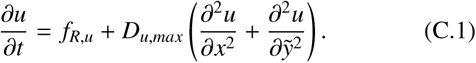

If we enforce the same scaling to 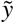 on all other equations, we get for every component *w* except *u*:

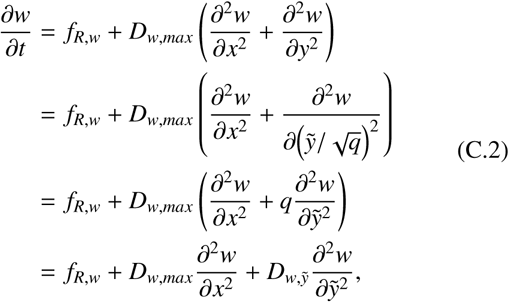

where 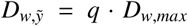. This means that reducing vertical active ROP diffusion by a factor *q* is equivalent to compressing vertical space by a factor 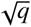, while raising the vertical diffusion of all other components by a factor *q*. This would suggest that a horizontal orientation can also be imposed on the final pattern by increasing the vertical diffusion of all components other than active ROP.

Selectively increasing the vertical diffusion of components may be difficult to achieve biologically. However, selectively reducing their horizontal diffusion seems intuitively equivalent and more realistically achievable in biology. We can demonstrate this as follows. Dividing equation C.2 by *q* yields:

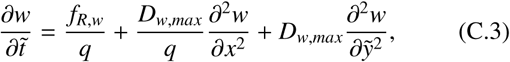

where 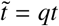. Transforming time in the same way for the equation of *u*, we obtain:

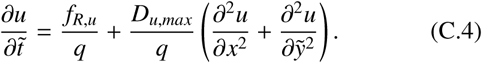

This means that vertical active ROP diffusion restriction is equivalent to horizontal diffusion restriction of all other components plus a scaling of *y* and *t*, a change in the parameters of the reaction function, and a reduction of the (isotropic) active ROP diffusion coefficient. Assuming the transformed parameters are chosen such that we remain in the Turing regime, we can therefore expect to obtain the same qualitative patterns, i.e., bands and spirals, for this alternative scenario.

Simulations show that horizontally restricting inactive ROP diffusion for the WPT model indeed leads to a horizontal orientation (Figure C.1), consistent with results from others [41]. While there are no biological indications for such a mechanism of imposing a horizontal orientation is involved in xylogenesis, it could be involved in other processes requiring banded patterns.

## D. Diffusion tensor rotation

For anisotropic diffusion in two dimensions, the diffusion term has the following form:

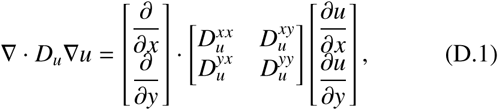

where 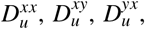 and 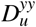 are the components of diffusion tensor *D*_*u*_, with 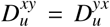. If diffusion is homogeneous, this simplifies to:

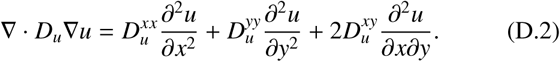

The components of the diffusion tensor can be derived from the diffusion rates in the unrestricted and restricted directions (1 and *ψ*, respectively) and the angle *ϕ* of the unrestricted direction with respect to the horizontal direction. For unrestricted diffusion along the *x*-axis and restricted diffusion along the *y*axis, we have:

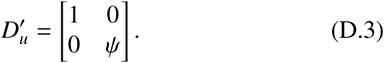

To obtain the diffusion tensor for the case where the unrestricted direction is rotated at an angle *ϕ* with respect to the x-axis, we need to rotate *D′*_*u*_ using rotation tensor *R*(*ϕ*):

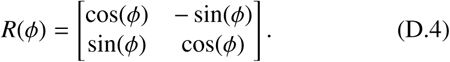

This way, we can obtain the diffusion tensor *D*_*u*_:

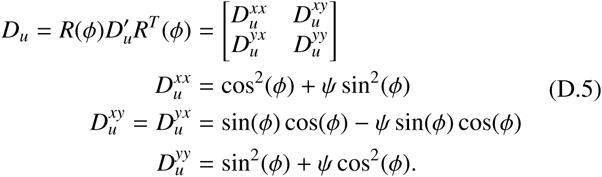

## E. Preferred spiral angles

With a strictly vertical reduction in the diffusion of active ROP, a fully horizontal banded pattern may form (Fig. 2). The distance between these bands is determined by the dynamics of the system and the zero flux boundaries at the top and bottom, which demand that the pattern starts and ends with either a peak or a valley. This would mean that the preferred number of bands based on the dynamics alone (*n*_*bands*_) should fall within a quarter of a band from the observed number of bands. However, the final steady state is also constrained by the initial emergence of the pattern, which depends on the random noise of the initial condition. Therefore, the final pattern may also contain a suboptimal number of bands, which increases the uncertainty on estimates of *n*_*bands*_. The only real ways around this flaw would be to solve the system on an infinitely large domain, which is not feasible, or to average over many simulations with different random seeds for the initial conditions, which is computationally expensive. Therefore, we make a basic estimate of *n*_*bands*_ and its uncertainty using the average, minimum and maximum numbers of bands observed in a limited number of repetitions.

Assuming for the moment that we know the preferred number of bands, we can calculate the preferred distance *d* between the bands as:

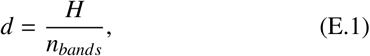

where *H* is the height of the domain. Assuming this distance is maintained in a spiral pattern, the angle *ϑ* of the spiral pattern depends on distance *d* and the vertical distance between spiral bands *d*_*y*_:

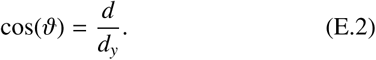

Vertical distance *d*_*y*_ is constrained by the periodicity of the spiral. The spiral covers a vertical distance *D*_*y*_ as it wraps around a domain of width *W*. This vertical distance *D*_*y*_ must be divisible by the vertical band-band distance *d*_*y*_, such that:
 

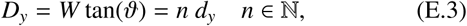

where *n* = 1 corresponds to a single spiral, *n* = 2 to a double spiral, etc. From equations E.1, E.2 and E.3, we can calculate the admissible angles of the spiral pattern if the distance between bands is determined solely by the system dynamics and not by boundary effects:

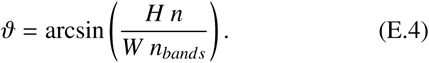

**Figure C.1:**
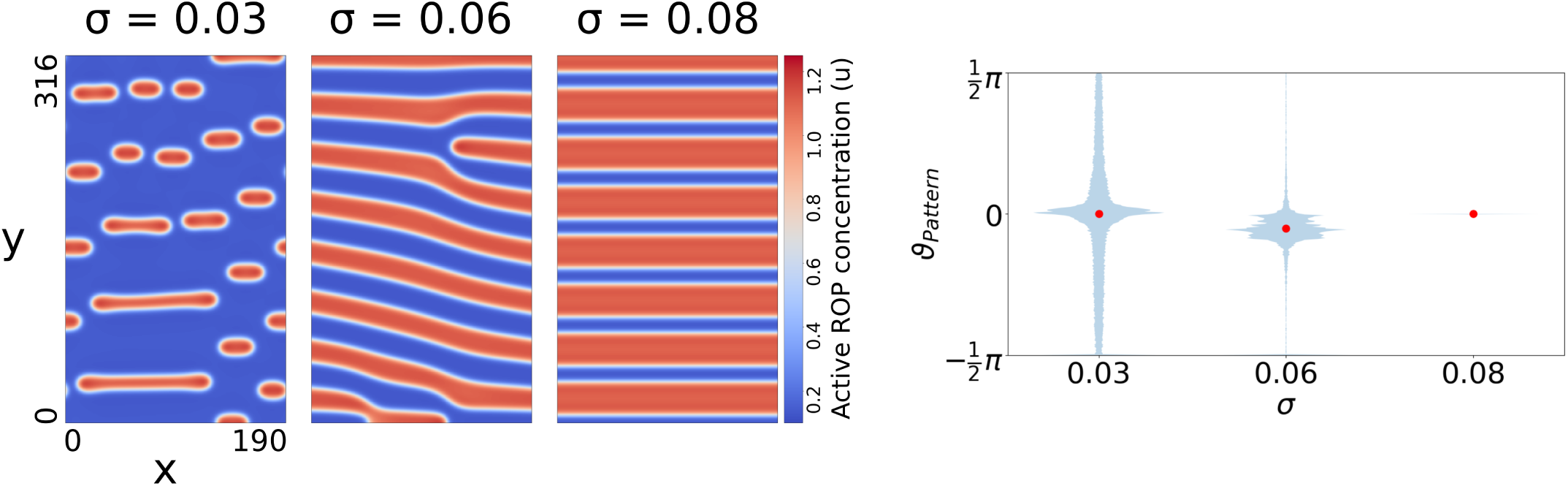
Horizontal diffusion restriction of inactive ROP can also impose a horizontal orientation on the pattern. Horizontal diffusion of inactive ROP was reduced by a factor 4. Snapshots were taken at *t* = 120000. Other parameters were as in Fig. 2.

Taking into account the fact that our estimate of *n*_*bands*_ may be off by a quarter and the possibility of a suboptimal number of bands, we can determine an expected range for each admissible angle, using the minimum and maximum number of bands observed in simulations with vertical diffusion restriction:

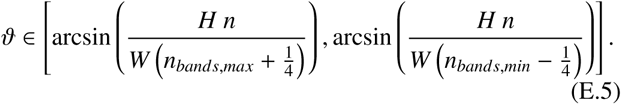

## F. Diffusion term decomposition

To study the influence of a pattern’s geometry on the effect of vertical diffusion restriction, we decompose the diffusion term in directions *z* and *w*. For any point on the domain, direction *z* runs in opposite direction of the gradient of *u*, so that derivative *u*_*z*_ = − |∇*u*|. Direction *w* runs perpendicular to *z*, so that *u*_*w*_ =0. To decompose the diffusion term, we ultimately need to express the second order spatial derivatives *u*_*xx*_ and *u*_*yy*_ in terms of *z* and *w*. The vectors 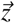 and 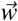 associated with these directions depend on the pattern angle *ϑ*, which we define as the angle between positive *x*-axis and the *w*-direction:

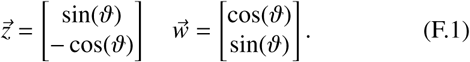

Using these vectors, we can express derivatives *u*_*z*_ and *u*_*w*_ in terms of *u*_*x*_ and *u*_*y*_:

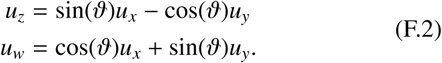

Solving for *u*_*x*_ and *u*_*y*_ gives:

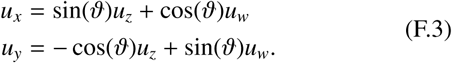

Repeating this process with *u*_*x*_ and *u*_*y*_ substituted for *u*, gives us *u*_*xx*_ and *u*_*yy*_:

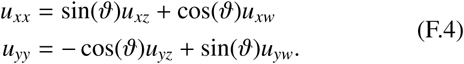

Substituting Eq. F.3 into Eq. F.4 gives:

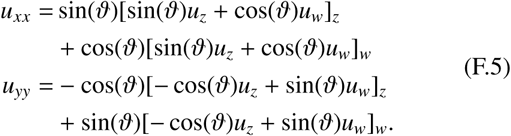

Since *u*_*w*_ = 0, we can simplify this to:

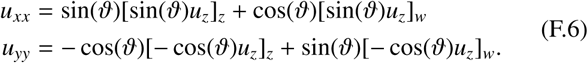

Applying the product and chain rules gives:

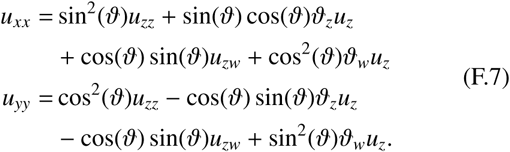

Since *z* is the direction of the gradient, the pattern angle does not change in this direction, so *ϑ*_*z*_ = 0. Also, *u*_*zw*_ = *u*_*wz*_ = [*u*_*w*_]_*z*_ = 0_*z*_ = 0. Therefore, the second and third terms of both equations vanish, leaving:

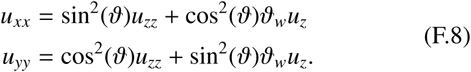

Substituting these derivatives into Eq. 5 gives:

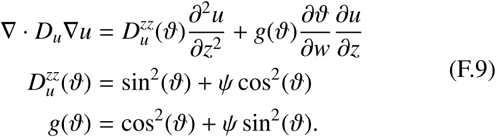

## G. Parameters of models with ROP-dependent ROP diffusion

Parameters used for the functions linking ROP activity and ROP diffusion are given in Table 1. For the instantaneous ROP effect (Fig. 7B), the maximally restricted diffusion coefficient (*ψ*_*min*_) was set to 0.25 and other parameters were chosen such that the final value of *ψ* would be close to its minimum for low ROP activity and close to its maximum for high ROP activity. For the simple delayed variant (Fig. 7C–E), the same value of *ψ*_*min*_ was taken and the initial (homogeneous) microtubule density *ρ*_0_ was set such that vertical ROP diffusion would start at its minimum (*ψ* = *ψ*_*min*_). For the model with positive feedback in microtubule density (Fig. 8), we tuned the parameters of the density equation to get a bistability in microtuble density that could be switched at low or high ROP levels. We used the same initial microtubule density (*ρ*_0_ = 2) for all simulations. To keep the starting conditions of the vertical active ROP diffusion coefficient close to that of used for the simple delay, *ψ*_*min*_ was lowered to 0.1, with *K*_*D*_ = 1. To keep vertical ROP diffusion close to its maximum at low microtubule densities, hill exponent *n*_*D*_ was set to 2, creating a sigmoidal curve.

## Notes

#### Summary of Updates

Small changes; new supplementary figure; extra appendix

