## Supplementary figures for "Robust banded protoxylem pattern formation through microtubule-based directional ROP diffusion restriction"

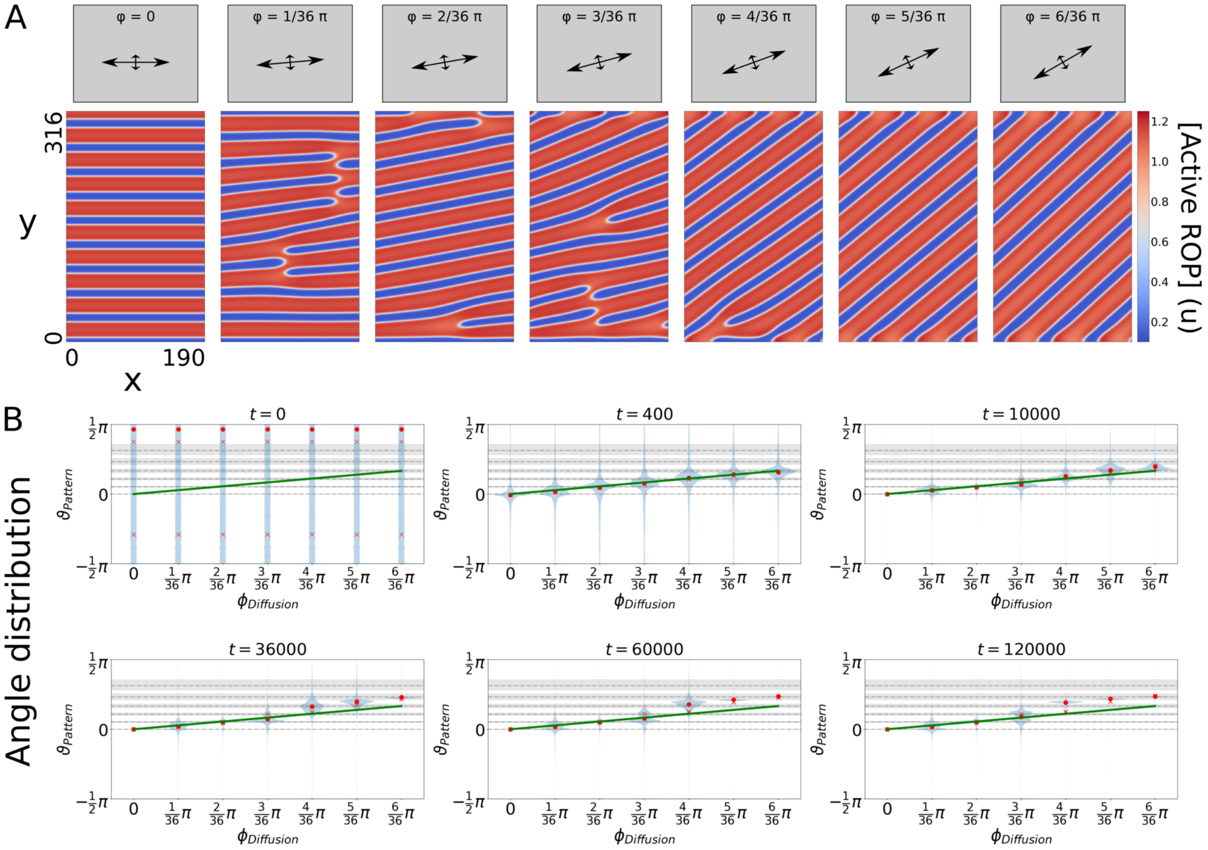


**Figure S.1:** Oblique diffusion restriction of active ROP results in spirals with angles $\vartheta$ depending on the diffusion angle $\phi$. (A) Active ROP concentrations at $t = 120000$ (close to steady state) in the gap regime of the unrestricted WPT model ($\sigma = 0.08$) for $\psi = 0.25$. (B) Distributions of the pattern angles (defined as orientation perpendicular to the gradient) are shown for various time points. Circles indicate the average orientation, crosses indicate average orientations for replicate runs with different initial conditions (for most time points crosses are invisible due to overlap with circles), solid green lines indicate the angle of maximum diffusion $\phi$, dashed lines indicate estimated angles for spiral patterns with band-band distances as in the banded patterns of Fig. 2, and shaded regions indicate uncertainty on those estimates due to boundary effects. Time-lapse videos of the corresponding simulations are available online.

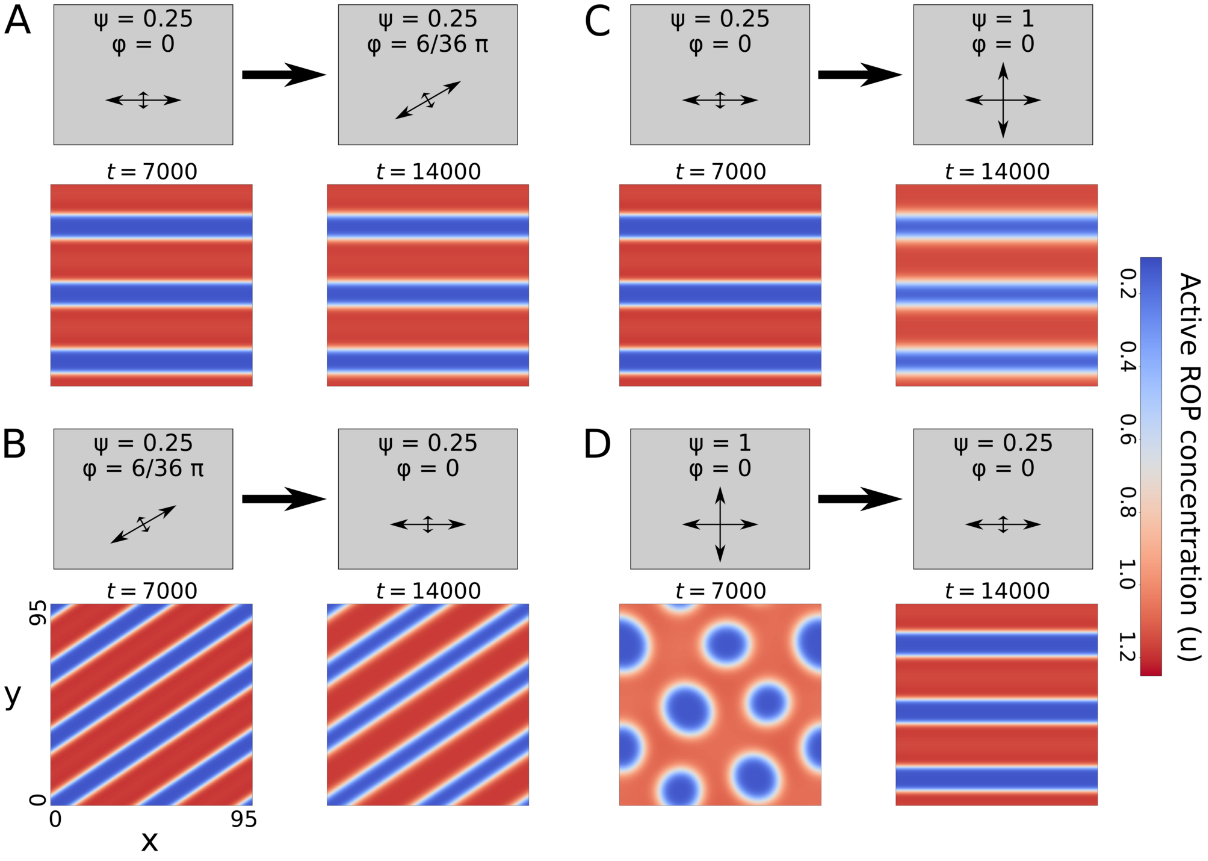


**Figure S.2:** Effect of altering diffusion restriction for established patterns of the WPT model in the gap regime ($\sigma = 0.08$). (A) Oblique diffusion restriction on horizontal bands. (B) Horizontal diffusion restriction on oblique bands. (C) Isotropic diffusion with horizontal bands. (D) Horizontal diffusion restriction on a pattern resulting from isotropic diffusion. Time-lapse videos of the corresponding simulations are available online.

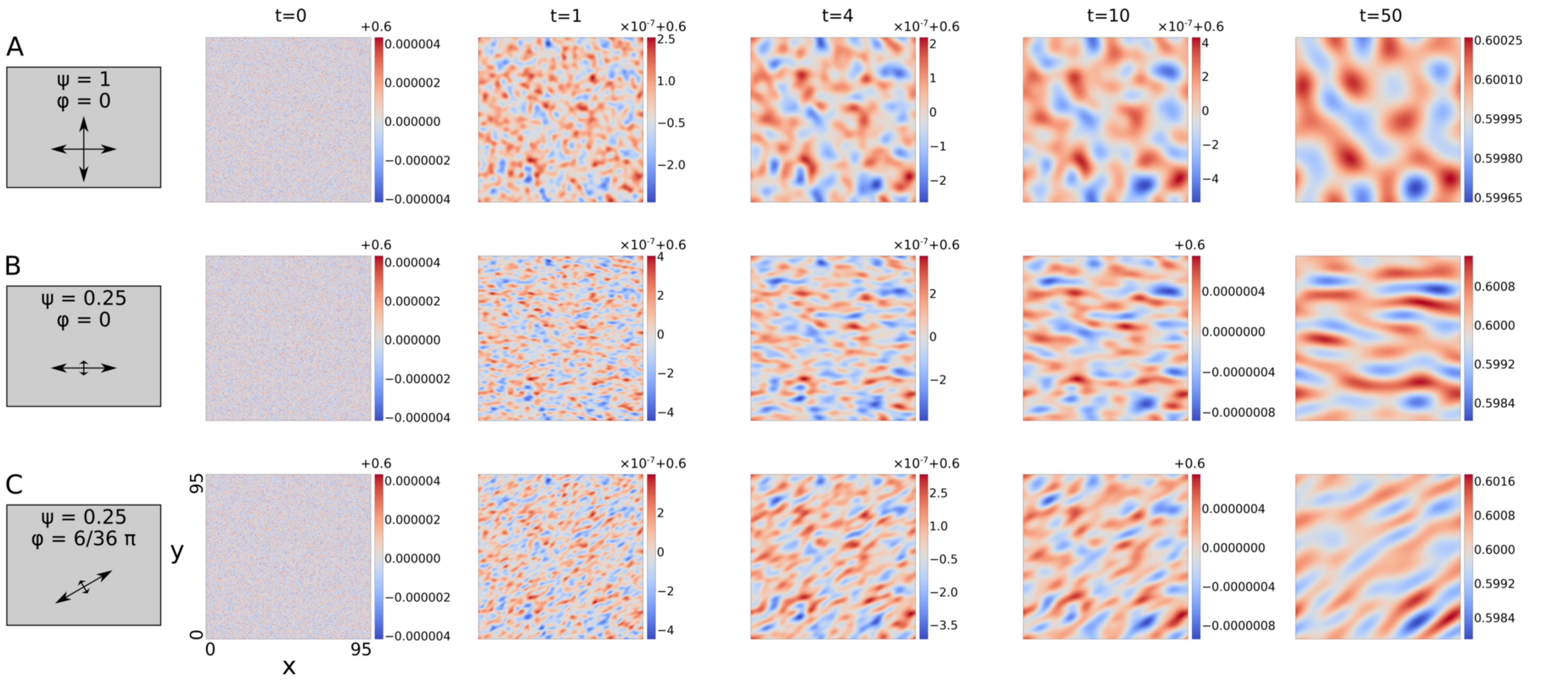


**Figure S.3:** Early stages of pattern formation with the WPT model ($\sigma = 0.06$). A. Isotropic diffusion. B. Horizontal diffusion restriction. C. Oblique diffusion restriction. Time-lapse videos of the corresponding simulations are available online.

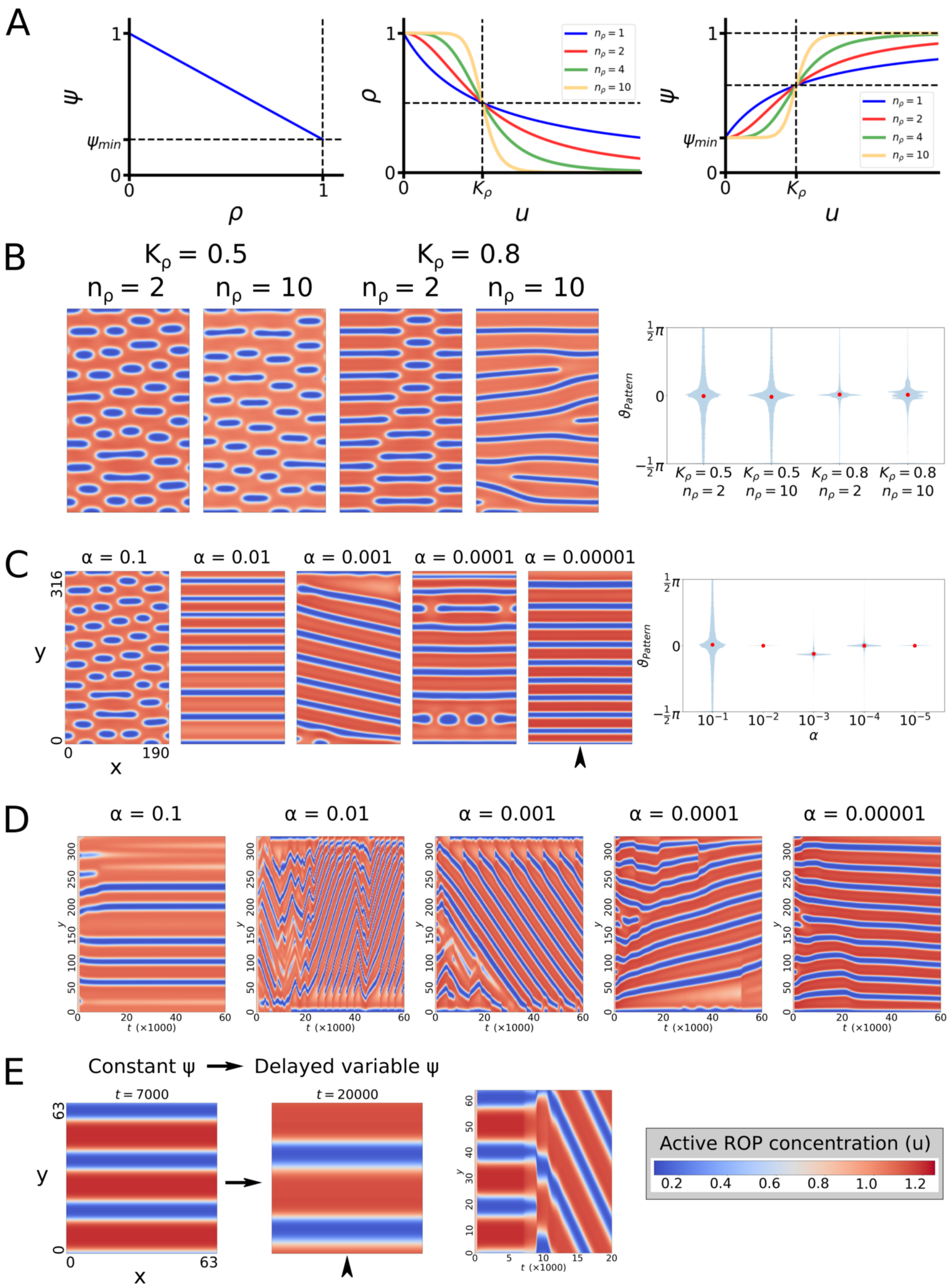


**Figure S.4:** Effect of ROP-dependent microtubule density reductions on the formation of banded patterns in the gap regime ($\sigma = 0.08$). (A) Relations between vertical diffusion component, (steady state) microtubule density, and active ROP concentration. (B) An instantaneous effect of ROP activity on microtubule density greatly hampers the robustness of the orientation mechanism. Snapshots at $t = 30000$. (C) Greater delays in the negative effect of active ROP on its own vertical diffusion restriction (smaller $\alpha$) result in straighter band formation. Snapshots at $t = 60000$. (D) Time evolution of concentrations from simulations shown in (C) at the horizontal position indicated by the arrowhead. Travelling waves (recognisable by diagonal lines) occur for larger delays (smaller $\alpha$), with wave speeds reducing as the delay becomes larger. (E) Travelling waves occur even when a static banded pattern is created first with a constant $\psi$ and delayed density reduction (delayed variable $\psi$) is initiated afterwards (at $t = 7000$, using $\alpha = 0.001$). Time evolution graph shows concentrations at the horizontal position of the arrowhead. Time-lapse videos of the corresponding simulations are available online.

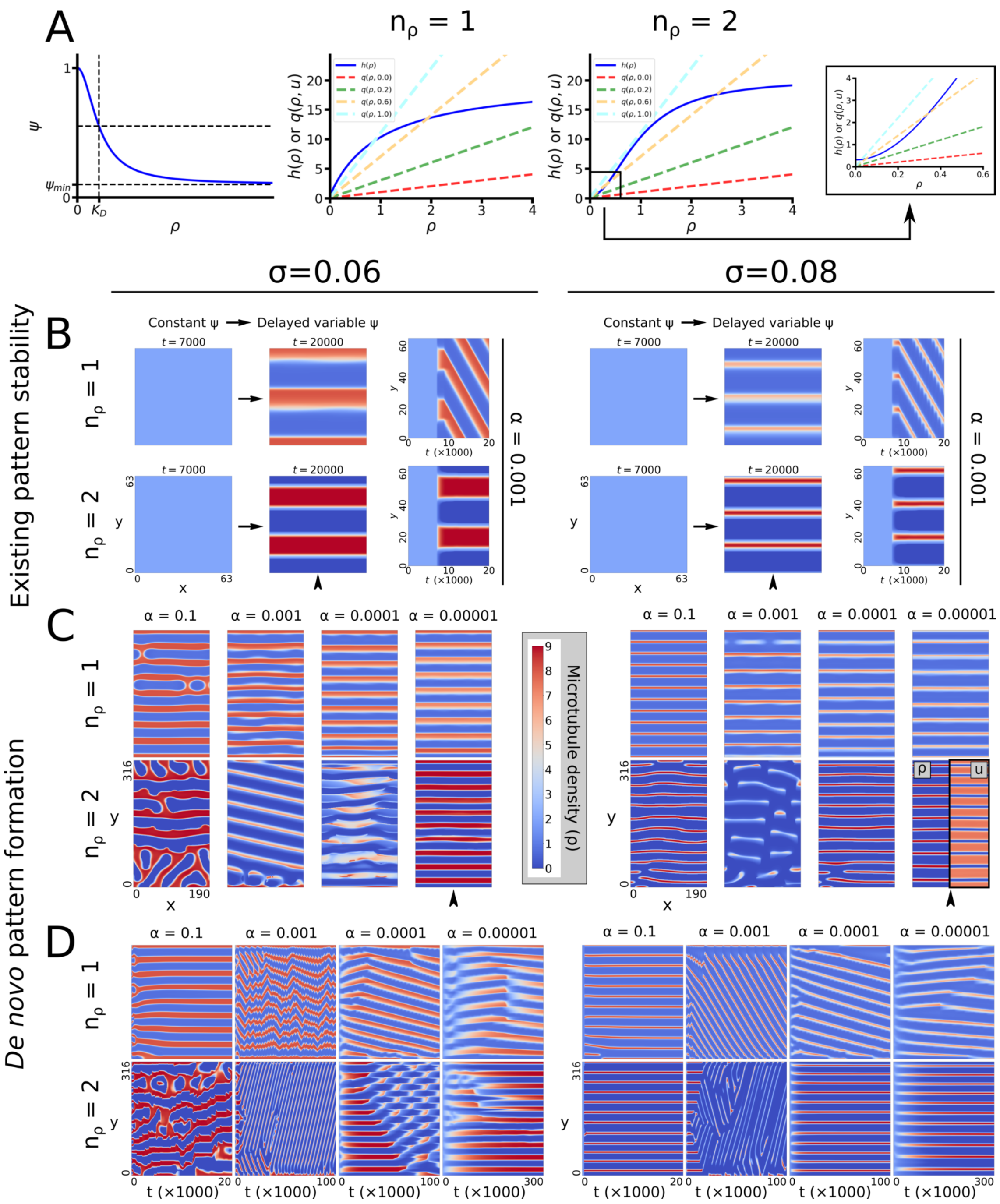


**Figure S.5:** Positive feedback in microtubule density can prevent bands from moving vertically. All simulations were performed in both the stripe ($\sigma = 0.06$) and the gap ($\sigma = 0.08$) regime of the WPT model. (A) Left: Relation between microtubule density ($\rho$) and vertical diffusion of active ROP. Right: Values of hill function and linear components of the microtubule density equation for various values of $u$. Intersections indicate steady states. Bistability can occur only for $n_\rho > 1$. (B) Stability of a banded pattern established with a constant microtubule density until $t = 7000$ for unistable ($n_\rho = 1$) and bistable ($n_\rho = 2$) microtubule dynamics. In all cases, $\alpha = 0.001$ and fully periodic boundary conditions were used. Graphs to the right show time evolution of microtubule densities at the horizontal position of the arrowhead. (C) Full *de novo* patterning with positive feedback of microtubule density starting from the homogeneous state for ROP concentrations and $\rho_0 = 2$. Snapshots were taken at the final time points indicated in (D). Inset in bottom right snapshot shows active ROP concentrations ($u$) for comparison. (D) Time evolution of simulations from (C) at the horizontal position indicated by the arrowhead. Corresponding active ROP concentrations for the entire figure are shown in Fig. 8 in the main text.
